# Systematic analysis of lectin gene family reveals dynamic modes of paralogue evolution and immune regulatory functions in tomato

**DOI:** 10.1101/2025.07.29.667230

**Authors:** Vishnu Shukla, Abin Panackal George, Rama Sai Venkata Marthi, Abhilasha Prashant Sonawane, Sanskruti Parida, Eswarayya Ramireddy

**Author notes:** **Corresponding author: Dr. Eswarayya Ramireddy**, Associate Professor, Biology Division, Indian Institute of Science and Education Research (IISER) Tirupati, Srinivasapuram, Jangalapalli Village, Panguru (G.P), Yerpedu Mandal, Tirupati-517619, Andhra Pradesh, India.

## Abstract

Lectins are a structurally diverse class of carbohydrate-binding proteins implicated in plant development and stress responses. In this study, we present a comprehensive structural, evolutionary, and functional analysis of the lectin gene family in tomato (*Solanum lycopersicum* cv. Heinz). We identified 247 lectin genes across eight major families, exhibiting diverse domain architectures, including merolectins, hololectins, and chimerolectins. Gene ontology enrichment, tissue-specific expression profiling, and stress-responsive transcriptomics highlighted key roles of lectin genes in development and pathogen defense. Comparative genomic analyses across five *Solanum* species and ancestral outgroups (*Vitis vinifera* and *Amborella trichopoda*) revealed both conserved and lineage-specific expansions driven by whole-genome and small-scale duplications. Evolutionary modeling indicated that most lectin gene duplications are under purifying selection, with some genes exhibit signatures of adaptive evolution. Expression and co-expression variance among duplicated paralogues revealed three distinct evolutionary fates -balance, dominance, and divergence. Notably, small-scale duplications frequently led to functional divergence, whereas whole-genome duplicates largely retained ancestral expression patterns. Functional validation through virus-induced gene silencing (VIGS) of two GNA chimerolectins, *Solyc04g077390* and *Solyc07g063700*, revealed their role as negative regulators of immunity. Silencing these genes enhanced resistance to *Ralstonia solanacearum*, reduced bacterial colonization, and triggered stronger induction of defense-related genes. Together, these findings highlight the structural innovation, evolutionary dynamics, and immune regulatory roles of lectin genes in tomato, and identifies promising targets for breeding disease-resistant cultivars in *Solanum lycopersicum* and related crops.

## Introduction

Plant lectins are carbohydrate-binding proteins that specifically and reversibly recognize distinct sugar moieties both within the plant and from external sources. These proteins are ubiquitously expressed across diverse plant tissues and play crucial roles in storage, signaling, development, and immune responses (Nonomura and Benson, 2014; Van Damme, 2022). Based on their carbohydrate-recognition domains (CRDs), plant lectins are classified into 12 families: *Agaricus bisporus* agglutinin, Amaranthin, class V chitinase-related (CRA), Cyanovirin, *Euonymus europaeus* lectin (EUL), *Galanthus nivalis* agglutinin (GNA), Hevein, Jacalin-related lectin (JRL), Legume lectin, LysM, *Nicotiana tabacum* agglutinin (Nictaba), and Ricin B lectin families (Eggermont et al., 2017). Structurally, lectins are further classified into four types based on domain organization: merolectins (single lectin domain), hololectins (two or more identical lectin domains), chimerolectins (a lectin domain fused with non-lectin domains), and superlectins (two or more distinct lectin domains) (Tsaneva and Van Damme, 2020).

Advances in genome and transcriptome sequencing have enabled comprehensive identification of lectin gene families—or “lectomes”—across various plant species. In *Arabidopsis thaliana*, 217 lectin genes spanning nine families were identified, with the majority encoding chimeric proteins involved in stress and immune responses (Eggermont et al., 2017). Similar genome-wide analyses in *Citrus sinensis*, *Cucumis sativus*, *Morus notabilis*, and *Glycine max* identified 141, 146, 197, and 359 putative lectin genes, respectively, often featuring conserved sequences and cis-regulatory elements linked to biotic and abiotic stress responses (Dang and Van Damme, 2015; Van Holle and Van Damme, 2015; Saeed et al., 2016; Ahmed et al., 2023). Many of these lectins harbor additional domains such as glycoside hydrolases (GHs), protein kinases, and F-box motifs, indicating multifunctionality in signaling and defense.

Owing to their critical roles in stress mitigation, lectins have been the focus of numerous functional studies. For example, *OSJAC1*, a monocot chimeric jacalin, confers broad-spectrum resistance to pathogens and enhances abiotic stress tolerance in rice and other crops (Weidenbach et al., 2016; Esch et al., 2012). Beyond defense, lectins are also involved in symbiotic interactions. *PtLecRLK1*, a lectin receptor-like kinase (LecRLK), mediates root symbiosis in *Populus* and improves symbiotic performance when overexpressed in *Arabidopsis* and switchgrass (Labbé et al., 2019; Qiao et al., 2021). LecRLKs, particularly from the GNA family, have been implicated in pathogen recognition and immune signaling (Yamaji et al., 2012; Luo et al., 2017). Interestingly, certain GNA-LecRLKs may act as susceptibility factors; for instance, *ERN1* in *Arabidopsis* increases vulnerability to root-knot nematodes (Zhou, et al 2023), suggesting that pathogens might co-opt lectin-mediated pathways originally evolved for beneficial symbiosis.

The evolution of plant lectins is characterized by gene family expansion and diversification in domain architecture from algae to higher plants. Many lectin families originated in charophyte algae and Streptophyta, with some ancestral architectures resembling animal lectins that were lost during land plant evolution, while others, such as the F-box/Nictaba combination, remain conserved in modern angiosperms (Van Holle and Van Damme, 2019). The expansion of lectin genes has also been associated with increased ploidy levels (Naithani et al., 2021). A broad survey across angiosperms demonstrated that gene duplicability is non-random: single-copy genes tend to regulate core cellular functions, while multi-copy genes are enriched in environmentally responsive pathways like signaling, metabolism, and transport (Li et al., 2016). Lectin genes fall into the latter category, as supported by their extensive duplication in higher plants (Van Holle et al., 2017). Gene family expansions—particularly in legume L-type LecRLKs and the Nictaba family in soybean—are attributed to both whole-genome and tandem duplication events (Hofberger et al., 2017; Van Holle et al., 2017). However, the relationship between lectin gene duplicability, domain evolution, and function remains poorly understood.

Despite growing evidence on the expansion and functional roles of lectin gene families in various plant species, comprehensive evolutionary and functional studies in tomato remain limited. Prior analyses have focused narrowly on specific lectin types or receptor kinases (Osman et al., 2024), without integrating domain architecture, duplication history, and expression dynamics across cultivated and wild tomato species. In this study, we performed a comprehensive comparative analysis of lectin gene families in *Solanum lycopersicum* (cv. Heinz), its wild relatives, and ancestral species. Our approach combined domain-based classification, phylogenetic reconstruction, orthology modeling, and expression dynamics to dissect the structural, evolutionary, and transcriptional diversification of lectin genes. We further functionally validated two GNA chimerolectins using virus-induced gene silencing (VIGS), demonstrating their role as negative regulators in tomato’s defense against *Ralstonia solanacearum*. These findings advance our understanding of lectin gene evolution and provide promising targets for enhancing disease resistance in tomato and related Solanaceae crops.

## Materials and methods

### Computational identification, functional annotation and expression profiling of lectin gene family

#### Data resource

The genomic datasets of five Solanum genomes (*S. lycopersicoides, S. pennelli, S. pimpinellifolium*, *S. lycopersicum var. Cerasiforme* and *S. lycopersicum cv. Heinz*) was downloaded from Sol Genomics Network (https://solgenomics.net/). Ensembl Genomes 53 (https://plants.ensembl.org/) was utilized to download genomic datasets of *V. vinifera* (PN40024.v4) and *A. trichopoda* (AMTR1.0). The hidden markov models (HMMs) profile of the lectin domains (*Agaricus bisporus* agglutinin; ABA - Pfam: PF07367, *Amaranthus caudatus* agglutinin; amaranthin - Pfam: PF07468, *Nostoc ellipsosporum* agglutinin; cyanovirin - P81180.2, Pfam: PF08881, *Euonymus europaeus* agglutinin; EUL - Pfam: PF14200, *Galanthus nivalis* agglutinin; GNA - Pfam: PF01453, *Artocarpus integer* agglutinin; JRL - Pfam: PF01419, *Glycine max* agglutinin; Legume - Pfam: PF00139, *Brassica juncea* agglutinin; LysM - Pfam: PF01476, *Hevea brasiliensis* agglutinin; Hevein - Pfam: PF00187, *Robinia pseudoacacia* agglutinin; CRA - Pfam: PF00704, *Nicotiana tabacum* agglutinin; Nictaba - Pfam: PF14299, and *Ricinus communis* agglutinin; Ricin-B - Pfam: PF00652) was downloaded from Pfam 35.0 (https://pfam.xfam.org/).

#### Identification and GO enrichment analysis

The putative lectin family genes in *V. vinifera*, *A. trichopoda,* and five Solanum genomes were identified using HMMER v3.2.1 (https://hmmer.org/) using ‘hmmsearch’ module with ‘trusted cut-off’ as the threshold. The resulting lectin proteins with E-value ≤ 1e-05 were queried to construct specific profile HMMs for each of the seven target genomes using the ‘hmmbuild’ module. These new species-specific profile HMMs were then used to search the corresponding lectin protein sequences in seven target genomes (Table S1). The lectin domain patterns in each putative protein sequence identified in seven genomes was validated using NCBI batch CD-search (Wang et al., 2023) and compared to each other (Table S2). The gene ontology (GO) enrichment analysis of lectin genes was performed using Shiny GO 0.77 (http://bioinformatics.sdstate.edu/go/) with FDR cut-off 0.05 (Table S3).

#### Transcription factor (TF) binding site enrichment analysis

For TF binding site enrichment, the 2kb promoter sequence of lectin genes from each family was retrieved from *S. lycopersicum cv. Heinz1702* genome (ITAG4.0) and scanned using MEME suite AME web tool (McLeay and Bailey, 2010) for TF binding sites by mapping on two plant-specific databases (JASPAR CORE plants and Arabidopsis) with default parameters except, the *e*-value report threshold was changed to 2000. The binding site enrichment of TF family members in promoters of each lectin family was calculated by comparing the total enrichment of corresponding TF family binding sites against the entire tomato genome (Table S4).

#### Meta-transcriptome expression analysis

Transcriptomic data comprising 22 tissue-specific and 19 stress-responsive datasets for *Solanum lycopersicum* lectin genes were retrieved from NCBI and the Plant Gene Expression Omnibus (PEO). For tissue-specific expression, TPM values were obtained directly from PEO, and datasets related to stress treatments were excluded. Where TPM was not provided, it was calculated using read counts and gene lengths (extracted from the ITAG4.0 GFF file via Galaxy) with the formulas: RPKM = Read Count / Gene Length; TPM = (RPKM / ΣRPKM) × 10C. TPMs were further extracted and scaled using R (Table S5). For tissue-specific expression analysis, expression values from all datasets were utilized to compute the tissue specificity index (Tau) for each lectin gene across various developmental tissue types. The calculation followed the widely recognized formula proposed by Yanai et al. (2005): τ=i = 1n(1-x_i_/x_max_)/n-1 where x*_i_* represents the expression level in tissue *i*, x*_max_* is the maximum expression across all tissues for the gene, and *n* is the total number of tissues (Table S6). For stress-responsive expression, curated NCBI datasets with available expression (log_2_FC) values were used directly. In datasets with only read counts, log_2_FC was calculated using DESeq2 (Table S7). Lectin-specific values were extracted, and heatmaps were generated using R.

### Comparative genomics analysis

#### Synteny survey

For establishing an intra-genomic syntenic relationship within each of the Solanum genomes, collinear lectin gene pairs were identified using MCScanX (Wang et al., 2024) with default parameters (Table S8). Following collinearity, the ‘duplicate_gene_classifier’ utility of MCScanX was used to classify lectin genes into different duplication categories including singleton, tandem, proximal, dispersed and segmental. To establish inter-genomic relationship between *V. vinifera* and five Solanum genomes, orthologous gene groups were identified using Orthofinder (Emms and Kelly, 2019) (Table S9). Taking protein sequences from all the genomes based on HMMER scan (Table S1) statistics, an all-versus-all sequence similarity search was performed using DIAMOND with an E-value threshold of 1e-5. Orthologous relationships, one-to-one, one-to-many and many-to-many, were inferred by reconciling gene trees with species tree (Table S10).

#### Phylogenetic analysis

The orthologous relationship was further validated using phylogenetic analysis between lectin sequences among *V. vinifera*, *Arabidopsis thaliana,* and five Solanum genomes. Multiple sequence alignments of 2009 protein sequences from each of the lectin family were synthesized using MAFFT version 7 (Katoh et al., 2019) using FFT-NS-i as an iterative refinement method with a gap opening penalty of 2.5. The alignment was trimmed using trimAl to remove columns that had more than 75% gaps (Capella-Gutierrez et al., 2009). IQ-TREE2 (minh et al., 2020) was used to infer the maximum likelihood tree based on protein sequence alignments. The JTT + G4 substitution model was applied, based on testing of 168 protein models using IQ-TREE2’s ModelFinder. Node support was assessed using 1,000 ultrafast bootstrap replicates and 1,000 SH-aLRT replicates. Similar analysis was performed for the phylogenetic tree construction of expansion and contraction subset of lectin genes except the substitution model used was JTT+F+G4 according to the ModelFinder. Both the trees were rooted using specified singleton lectin sequences from *A. trichopoda* and the final tree was visualized and annotated using iTOL (https://itol.embl.de/).

#### Gene family expansion/contraction and Ka/Ks estimation

Gene family expansions and contractions were analyzed using CAFE5 (Mendes et al., 2021), based on the ultrametric species tree and gene family count matrix generated by OrthoFinder (Table S11). CAFE5 was executed with a gamma-distributed rate model and the parameter *k* = 3 to account for variation in gene gain and loss rates across families and to detect lineage-specific changes in gene family size along the species tree. The resulting orthogroups with *P*-value less than 0.01 were considered as ‘expansion orthogroups’ or ‘contraction orthogroups’. The rates of sequence evolution (Ka/Ks) for both cross-species and paralogous lectin pairs were estimated using KaKs_Calculator v2.0 (Wang et al., 2010). Here, the input AXT files containing codon-aligned nucleotide sequences of gene pairs were trimmed and cleaned to ensure the preservation of correct codon structure.

### Expression-based paralogue evolution

#### Overview

Paralogous gene pairs were identified and considered for downstream analyses based on the integrated outputs from MCScanX, OrthoFinder, and CAFE5. MCScanX was used to detect collinear gene pairs and classify them into different duplication types. OrthoFinder was employed to define orthogroups and distinguish paralogues within and across species based on gene tree reconciliation. Additionally, gene families showing evidence of expansion or contraction were identified using CAFE5, which helped categorize paralogous pairs likely to have undergone conserved or lineage-specific diversification (Table S12). Only those gene pairs that were consistently identified as paralogs across these approaches were retained for further expression and evolutionary analyses. In total, we obtained 166 paralogue lectin pairs in *S. lycopersicum*, representing all possible gene pairs within expansion orthogroups. To investigate different modes of paralogue functional evolution, the transcript abundances and co-expression of paralogue pairs in each expansion orthogroup was compared in different tissues (apices, cotyledons, hypocotyl, flowers, inflorescences, fruits and roots) of *S. lycopersicum* and four other *Solanum* relatives (Benoit et al., 2025).

#### Co-expression and expression analysis

Briefly, the expression pattern of paralogue pairs of *S. lycopersicum* lectins (Table S13) in each expansion orthogroup was compared with the expression patterns of their corresponding orthologs in other wild and domesticated *Solanum* relatives namely, *Solanum macrocarpon* (Table S14), *Solanum candidum* (Table S15), *Solanum quitoense* (Table S16) and *Solanum prinophyllum* (Table S17). The normalized TPM counts of lectin genes (from expansion orthogroups) from each species was retrieved from datasets provided in Benoit et al., 2025 and orthologous genes between *S. lycopersicum* and other *Solanum* relatives were identified using reciprocal BLAST analysis. Coexpression networks for each species were constructed by computing Pearson correlation coefficients between all paralogue pairs. For each paralogue pair, the correlation values were ranked, with missing values (NA) assigned the median rank. The ranked correlation matrix was then standardized by dividing each value by the maximum rank per gene to normalize coexpression strength across the network.

#### Different modes of expression-based evolution

Paralogous gene pairs within each species were grouped into different retention categories based on their expression differences and coexpression across tissues (Benoit et al., 2025). These two factors, average fold-change in expression and coexpression, capture both the overall expression level differences and the similarity in expression patterns between paralogue pairs. Based on these metrics, the paralogue pairs were classified into four categories: **Balanced -** Higher coexpression (correlation > 0.8) with minimum differences in expression (mean logC[FC] < 1; standard deviation < 1). **Dominance -** Higher coexpression (correlation > 0.8), but one gene consistently shows higher expression (mean logC[FC] ≥ 1; standard deviation < 1). **Specialized -** Higher coexpression (correlation > 0.9) with both large and variable differences in expression (mean logC[FC] ≥ 1; standard deviation ≥ 1). **Diverged -** Weak coexpression (correlation < 0.4) and show both large and variable expression differences (mean logC[FC] ≥ 1; standard deviation ≥ 1).

### Bacterial wilt assay in tomato

#### Ralstonia solanacearum infection

The *R. solanacearum* F1C1-mCherry strain was obtained from S.K. Kumar Ray and cultured in BG broth (10 g peptone, 1 g yeast extract, 1 g casamino acid) at 28°C for 48 hours, as previously described by Kumar et al. (2017). To analyze the differential expression of candidate genes against *R. Solanacearum* infection, bacterial cells at a concentration of 1 × 10C CFU/mL were inoculated into the pots of two-week-old tomato plants using the drench method. The plants were maintained in a greenhouse at 28°C under a 16:8 h light/dark cycle with 80% humidity for five days, followed by RNA isolation and RTq-PCR.

For analyzing the phenotype of VIGS silenced plants, *R. solanacearum* of 1 × 10C CFU/mL were inoculated into the pots of 3-week-old tomato plants using the drench method. The plants were maintained in a greenhouse at 28°C under a 16:8 h light/dark cycle with 80% humidity. Disease symptoms were observed for seven days after inoculation and photographed using a DSLR camera. Each experiment included at least 12 infected plants and was independently replicated. The wilting index was determined following the method described by Hiles et al., 2024. Briefly, wilting symptoms were assessed using a scale from 0 to 4, where 0 indicated no visible symptoms,1 indicates ≤25% wilting symptoms, 2 indicates ≤50% wilting symptoms, 3 indicates ≤75% wilting symptoms and 4 represents ≤100% wilting of leaves. Based on these wilt scores, the disease index (DI) was calculated using the formula: DI = Nw/N (Nw = number of leaves wilted, N = total number of leaves).

#### VIGS-mediated silencing

The TRV based vectors pTRV1 and pTRV0 were obtained from Chandan, R. K., et 2023, NIPGR, New Delhi. The target region for VIGS silencing of *Solyc04g077390* and *Solyc07g063700* approximately 300 base pairs in length was predicted by SGN VIGS (http://vigs.solgenomics.net) (Table S18). The predicted regions were PCR amplified using specific target primer pairs and cloned into multiple cloning sites of pTRVO vector and confirmed by Sanger sequencing. The constructs were then transformed into *Agrobacterium tumefaciens* GV3101 cells. Subsequently the cultures of *pTRV:Solyc04g077390* and *pTRV:Solyc07g063700* were mixed with pTRV1 at 1:1 ratio(OD600∼0.1) and infiltrated into 2 week old tomato seedlings. *SlPDS* is used as positive control for Knock-down (Figure S7B). The silencing of *Solyc04g077390* and *Solyc07g063700* was confirmed using RT-qPCR (Figure S7C).

#### Colony Forming Unit assay (CFU assay)

The CFU of tomato-infected plants was quantified following the method described by Yu, Gang, et al., 2023. The shoot and root tissues of VIGS-*R. solanacearum*-infected plants were collected separately, and their fresh weights were recorded. The samples were then grounded in a 2 mL tube with 200 µL of sterile water using a Retsch mill at 30 cycles for 2 minutes, followed by dilution to a final volume of 1000 µL. Subsequently, 20 µL of the homogenate was added to 180 µL of sterile water, and serial dilutions were performed up to 10C. A 100 µL aliquot from the 10C dilution was spread onto BG agar plates containing 30 µg/mL gentamycin and incubated at 28°C. *R. solanacearum* colonies were observed after 36 hours, counted and quantified per milliliter. The experiment was performed in 6 biological replicates.

#### Xylem area measurement

The xylem area was measured in infected and control stem sections from the crown region of VIGS knockdown plants. Stem sections were prepared using a vibrating microtome (VF-310-OZ) and stained with 0.01% toluidine blue for 5 minutes. This was followed by dehydration through a graded ethanol series (20%, 50%, 70%, 90%, and 100%). The samples were then observed under a light microscope and imaged using a Leica DM2000 LED. The xylem vascular area was quantified using ImageJ software. The toluidine blue-stained xylem region was outlined using the color threshold and magic wand tool, and the area was measured using the “region of interest” function.

#### Validation of gene expression using RT-qPCR

The RNA-seq expression profiles of lectin genes during tomato development and in response to biotic stress were validated using RT-qPCR-based transcript analysis. Total RNA was extracted from three biological replicates of developmental tissue samples collected from 90-day-old plants, as well as from plants five days after *Ralstonia solanacearum* infection. RNA isolation was performed using the Takara RNA extraction kit (Cat. No. 6110A), following the manufacturer’s protocol. For cDNA synthesis, 2 µg of total RNA was used. Similarly, to quantify the transcript levels of candidate genes in knockdown plants and response genes in VIGS-*R. solanacearum*-infected plants, 1 µg of RNA from root and shoot was used for cDNA synthesis following the same protocol. The cDNA samples were used to determine transcript levels using RT-qPCR as described in Ramireddy et al 2018. The experiment was performed in three biological replicates (n = 3 seedlings per replicate). The primers used are listed in Table S18.

#### Statistical analysis

The wilting index was assessed using the Friedman and Duncan tests to evaluate disease progression within the lines, while the Wilcoxon signed-rank test was used for comparisons between the lines. For colony-forming unit (CFU) analysis, significance was determined using Student’s t-test. Similarly, xylem area significance was analyzed using a two-way ANOVA with Tukey’s HSD post hoc test. For relative expression analysis of response genes, a one-way ANOVA followed by Student-Newman-Keuls post hoc test was conducted. To analyze the differential expression of response genes and candidate genes upon *R. solanacerum* infection, salt stress and hypoxia Student’s t-test was performed.

## Results

### Structural and functional insights into tomato lectin gene family

Advancements in high-throughput genome sequencing have greatly facilitated the evolutionary and functional annotation of gene families in plants (Vaattovaara et al., 2019). In plants, 12 lectin gene families have been classified based on their carbohydrate recognition domains (CRDs) (Eggermont et al., 2017; Naithani et al., 2021). By querying all known CRDs in the tomato (*Solanum lycopersicum*, cultivar Heinz) genome, we identified 247 putative lectin genes (Table S1). Unrooted phylogenetic analysis, including orthologs from *Arabidopsis thaliana*, revealed that tomato lectin genes are distributed across eight major lectin families: GNA (95 genes), Legume (25), Malectin (44), Hevein (23), Nictaba (34), JRL (9), LysM (9), and CRA (8) (Figure 1A). Domain architecture analysis further classified these genes into three structural types: merolectins, hololectins, and chimerolectins (Figure 1B; Table S2). Chimerolectins were the most prevalent structural type; however, both merolectins and chimerolectins were present across all lectin families. In contrast, hololectins were confined to the JRL, GNA, and Hevein families. These results suggest that tomato lectin genes possess diverse domain organizations, likely contributing to a broad range of biological functions. The emergence of multi-domain proteins has been associated with increased genome complexity and functional diversification (Forslund et al., 2008). Notably, the expansion of receptor-like protein kinases in angiosperms is often linked to the fusion of lectin domains, particularly GNA and legume types (Xing et al., 2013). Previous studies have highlighted the role of complex lectins—such as chimeric, holo-, or super-lectins—in mediating responses to pathogenic signals and facilitating symbiotic interactions in legumes and cereals (Zipfel & Oldroyd, 2017). However, the presence of multiple domains alone does not confirm enhanced or novel functionality; this warrants further experimental validation.

**Figure 1.**
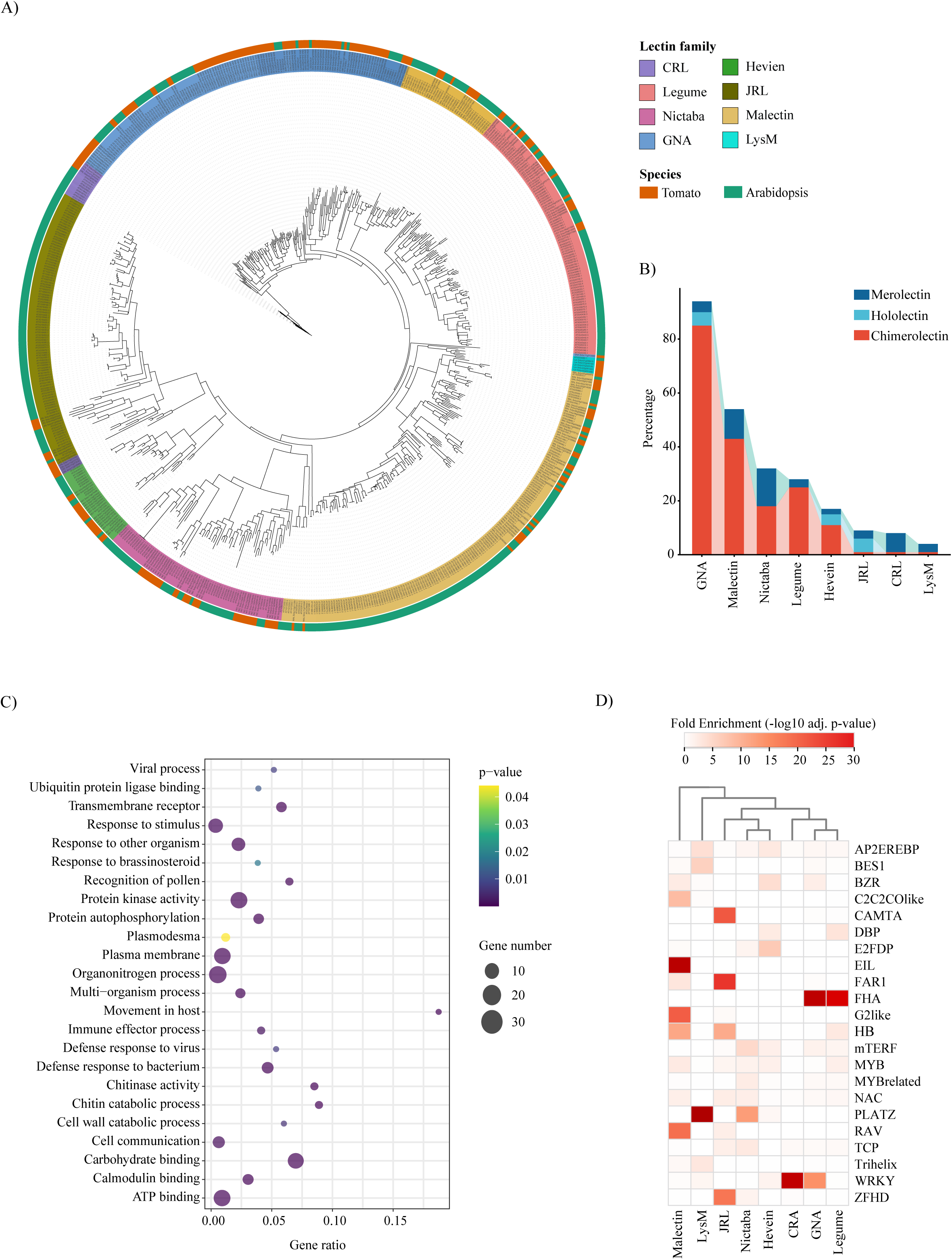
Functional annotation of the lectin gene family in the tomato genome. A) Maximum-likelihood phylogenetic tree showing the distribution of tomato lectin domain-containing proteins alongside Arabidopsis lectins. Distances are proportional to evolutionary distances and specified by scale bar (Tree scale: 10.) Both lectin gene families and species origins are color-coded for distinction. B) Filled stacked bar plot illustrating the distribution of various lectin types within each lectin family based on structural classification. The x-axis represents the percentage of lectin genes assigned to each structural category within a given family. C) Bubble plot illustrating Gene Ontology (GO) enrichment analysis of tomato lectin genes. The x-axis indicates the number of lectin genes associated with each GO term relative to the total number of genes annotated with that term in the tomato genome. Circle size corresponds to the number of lectin genes mapped to each GO term, and the color gradient reflects the statistical significance of GO term enrichment. D) Heatmap showing the enrichment of transcription factor (TF) binding sites in the promoter regions of lectin genes. The intensity of the color gradient represents the degree of enrichment for each TF family.

Gene Ontology (GO) enrichment analysis indicated that tomato lectin genes are significantly associated with biological processes such as defense response, protein phosphorylation, cell communication, and pollen recognition (Figure 1C; Table S3). To understand the transcriptional regulation of these genes, we analyzed the enrichment of transcription factor binding sites (TFBS) in their promoter regions (Figure 1D; Table S4). Distinct transcription factor (TF) families were associated with specific lectin subfamilies: FHA TF binding sites were enriched in GNA and Legume lectin promoters, while EIL, PLATZ, WRKY, and FAR1 motifs were overrepresented in the promoters of Malectin, LysM, CRA, and JRL genes, respectively.

Over the past two decades, numerous studies have highlighted the critical roles of lectins not only in mediating pathogen resistance (Willmann et al., 2011; Trontin et al., 2014; Wang et al., 2014), but also in regulating key aspects of plant development (Xin et al., 2009; Van Hove et al., 2015; Micol-Ponce et al., 2022). To further elucidate their functional significance in tomato, we investigated the tissue-specific expression patterns and stress-responsive transcriptional dynamics of lectin gene family members. Hierarchical clustering based on normalized TPM values revealed substantial variation in expression patterns across developmental stages (Figure S1; Table S5). Hevein family genes showed relatively high and widespread expression. For example, *Solyc10g055800* and *Solyc10g055810* were highly expressed in vegetative tissues, while *Solyc10g017970* and *Solyc10g017980* were upregulated during fruit and seed development. Beyond the Hevein family, several Malectin genes (*Solyc06g009540*, *Solyc10g054050*, *Solyc01g109950*, *Solyc01g108270*, *Solyc06g009550*, *Solyc11g010920*, *Solyc12g014350*) and Nictaba genes (*Solyc10g051160*, *Solyc10g083730*, *Solyc12g098200*, *Solyc07g006390*, *Solyc05g055870*, *Solyc01g091700*, *Solyc12g056410*) were specifically expressed in floral tissues, including young and mature flowers, ovules, and male gametophytic cells. Additionally, certain genes showed marked expression in specialized tissues—for instance, Hevein genes (*Solyc01g097280*, *Solyc10g074440*) in trichomes, and members of the Nictaba, Legume, and GNA families in the root apical meristem (*Solyc09g008820*, *Solyc09g007510*, *Solyc01g006590*, *Solyc04g015460*). Further, to enable a more robust assessment of family-level trends, we calculated the tissue-specificity index (τ) for each lectin gene across various organs (Figure 2A; Table S6). In root tissues, the CRL, GNA, Nictaba, and Hevein families exhibited high specificity (τ > 0.8) toward the root apical meristem, whereas JRL genes showed distinct expression in lateral roots. In floral tissues, GNA, Legume, and Hevein genes were preferentially expressed in pollen grains, mature flowers, and styles; CRL and Nictaba genes were enriched in the pistil; and Malectin genes were specific to floral buds. During fruit development, JRL, Legume, and LysM genes were predominantly expressed during early stages, while CRL, GNA, and Hevein genes were associated with later stages. These observations were validated by high transcript abundance and τ values of specific genes, such as *Solyc10g017970* and *Solyc07g009530* (Hevein) in 28 DPA fruit, and *Solyc12g011290* (Malectin) in floral buds (Figure 2B).

**Figure 2.**
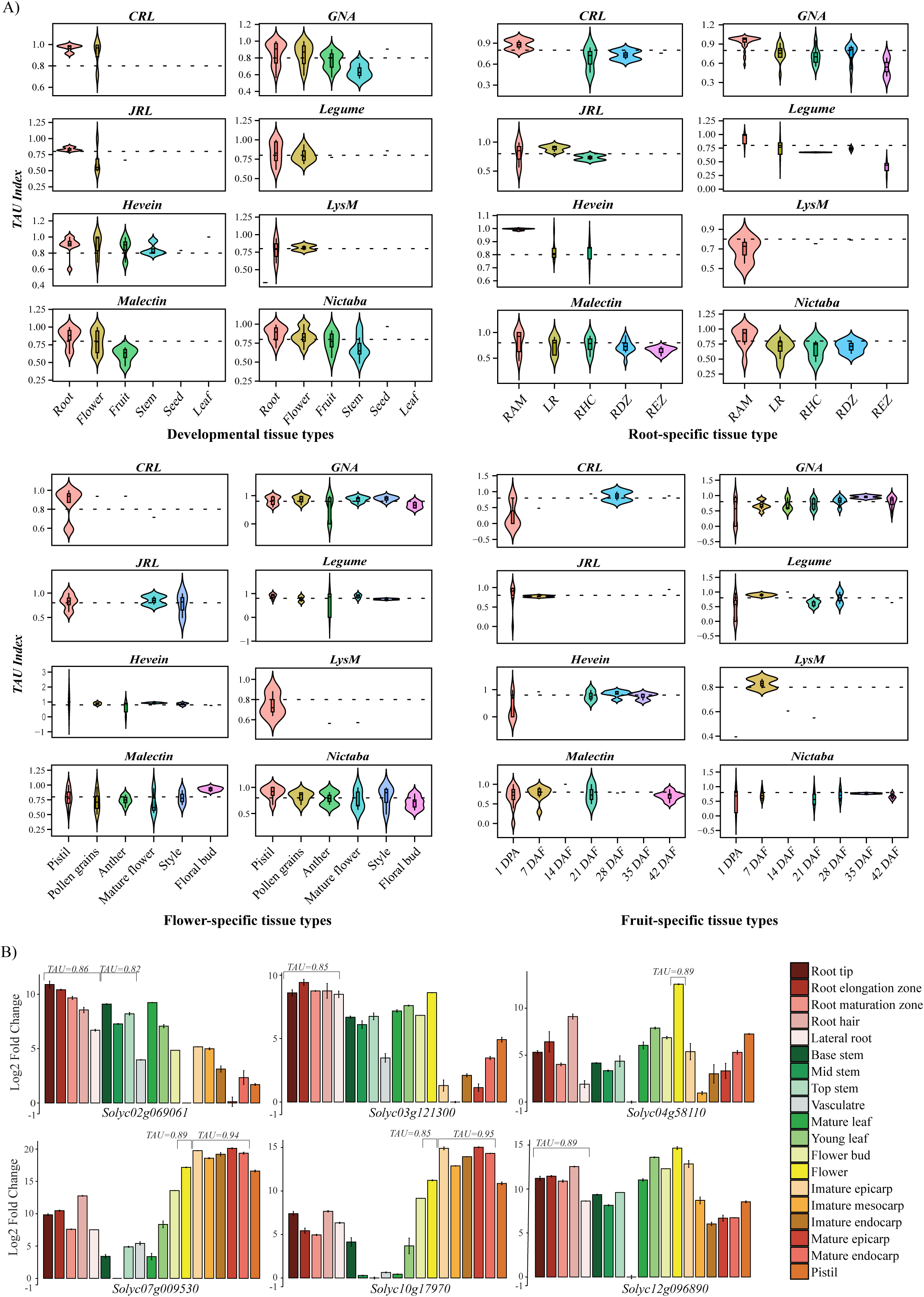
Tissue-specific expression of lectin gene families in *Solanum lycopersicum* (Heinz) during development. (A) Violin plots showing the distribution of Tau index values for different lectin gene families across various developmental tissues, as well as in root-(RAM, root apical meristem; LR, lateral root; RHC, root hair cell; RDZ, root differentiation zone; REZ, root elongation zone), flower-, and fruit-specific tissues (DPA, days post anthesis; DAF, days after fertilization). Each panel represents a specific tissue context, and gene families are grouped along the x-axis. Tau index values range from 0 (broad expression) to 1 (high tissue specificity). (B) Validation of tissue-specific transcript abundance of representative lectin genes by RT-qPCR across different developmental tissue types. The Tau index value for each gene is indicated above the bars of specific tissue type.

In addition to developmental expression patterns, we examined the transcriptional responses of lectin genes under various abiotic and biotic stress conditions (Figure S2; Table S7). Specific members of the LysM (*Solyc02g081040*) and JRL (*Solyc10g078600*) families exhibited significant downregulation under endoplasmic reticulum and salt stress, while a Nictaba gene (*Solyc02g068300*) showed reduced expression in response to both salt and cold stress. Conversely, a Malectin gene (*Solyc10g006870*) and a GNA gene (*Solyc07g063710*) were notably upregulated under drought conditions. Several Nictaba genes (*Solyc12g096890*, *Solyc01g100010*, *Solyc02g069055*) were consistently downregulated during heat stress, suggesting their potential roles in temperature sensitivity. Under biotic stress conditions, JRL gene (Solyc09g090445), Hevein genes (*Solyc07g009530*, *Solyc10g055810*) and a GNA gene (*Solyc02g076830*) were upregulated at different stages of anthracnose disease development, indicating their possible involvement in fungal pathogenesis. Several GNA genes (*Solyc10g005440*, *Solyc07g063700*, *Solyc02g079530*) were also found to be upregulated in response to bacterial infection such as *Pseudomonas syringae*, *Ralstonia solanacearum* and *Xanthomonas euvesicatoria*. Notably, a CRL gene (*Solyc07g005110*) showed highly specific induction in response to Potato Virus X (PVX), Tomato mosaic virus (TMV), and potato spindle tuber viroid, underscoring its potential role in broad-spectrum stress defense. These stress-responsive expression patterns reinforce the findings from GO enrichment analysis, which highlighted lectin genes’ involvement in critical biological processes such as “defense response” and “cell communication”. The transcriptional plasticity of specific lectin genes under diverse environmental challenges suggests that they play important roles in adaptive responses, possibly through signaling pathways mediated by their carbohydrate-binding functions.

### Evolutionary conservation and divergence of lectin genes in tomato

Comparative analysis of multiple plant genomes has been instrumental in uncovering the evolutionary dynamics of gene families (Soltis and Soltis, 2020). To investigate the impact of angiosperm evolution—particularly the eudicot-wide whole genome triplication (WGT) event— and tomato domestication on the diversification of lectin genes, we performed a comparative genomic study involving two ancestral species (*Amborella trichopoda* and *Vitis vinifera*), wild tomato relatives (*Solanum lycopersicoides*, *S. pennellii*, and *S. pimpinellifolium*), and domesticated tomato varieties (*S. lycopersicum* var. *cerasiforme* and *S. lycopersicum* cv. Heinz) *A. trichopoda* represents a basal angiosperm lineage that has not undergone recent or lineage-specific genome duplications (Albert et al., 2013), while *V. vinifera* is considered a close outgroup to Solanaceae and has not experienced additional genome duplication since the ancestral eudicot WGT event (Jaillon et al., 2007). Therefore, these two genomes serve as key evolutionary references for inferring the origin, expansion, and diversification of lectin gene families in Solanaceae, particularly in tomato.

#### Intra-species synteny analysis

Intra-genomic synteny analysis across four *Solanum* genomes (wild and cultivated tomatoes) revealed significant duplication events involving lectin genes (Figure 3A; Table S8). A total of 104 collinear gene pairs were identified, suggesting that duplication has significantly contributed to the expansion of the lectin gene family. Based on collinearity, these duplications were primarily segmental, proximal, or tandem in nature. Ka/Ks analysis of these duplicated pairs indicated that most gene pairs are under purifying selection (Ka/Ks < 1), consistent with evolutionary pressure to maintain their function (Figure 3B; Table S8). A smaller subset showed signs of positive selection (Ka/Ks > 1), implying adaptive divergence in specific gene pairs. In *S. lycopersicoides*, 9 gene pairs were under strong purifying selection (Ka/Ks < 0.5), 23 under relaxed purifying selection (0.5 < Ka/Ks < 1), and 4 gene pairs under positive selection, including Nictaba (*Solyd02g053150*:*Solyd02g061180*), CRL (*Solyd01g077660*:*Solyd03g067260*; *Solyd01g077710*:*Solyd05g066960*), and Malectin (*Solyd02g064110*:*Solyd03g051080*). In *S. pimpinellifolium*, 9 pairs were under strong, 8 under relaxed purifying selection, and 2 gene pairs, Nictaba (*SPI01g02302*:*SPI12g03885*) and Malectin (*SPI03g05384*:*SPI06g03899*) showed positive selection. In *S. lycopersicum* var. *cerasiforme*, 7, 11, and 2 gene pairs fell under strong, relaxed, and positive selection, respectively, with the latter involving GNA paralogs (*SLYcer02g04363*:*SLYcer10g00399*; *SLYcer07g05414*:*SLYcer10g00402*). Similarly, in *S. lycopersicum* cv. Heinz, 11 pairs showed strong purifying selection, 13 relaxed purifying selection, and 3 gene pairs GNA (*Solyc02g079700*:*Solyc07g063770*; *Solyc01g094830*:*Solyc03g025190*) and Malectin (*Solyc03g121230*:*Solyc06g062450*) were under positive selection. These results underscore both the evolutionary conservation and lineage-specific adaptive diversification of lectin genes within *Solanum* species. However, integrating cross-species orthology with expression and selection pressure data may provide a more comprehensive framework to understand the evolutionary trajectory and functional specialization of lectin genes in tomato and related species.

**Figure 3.**
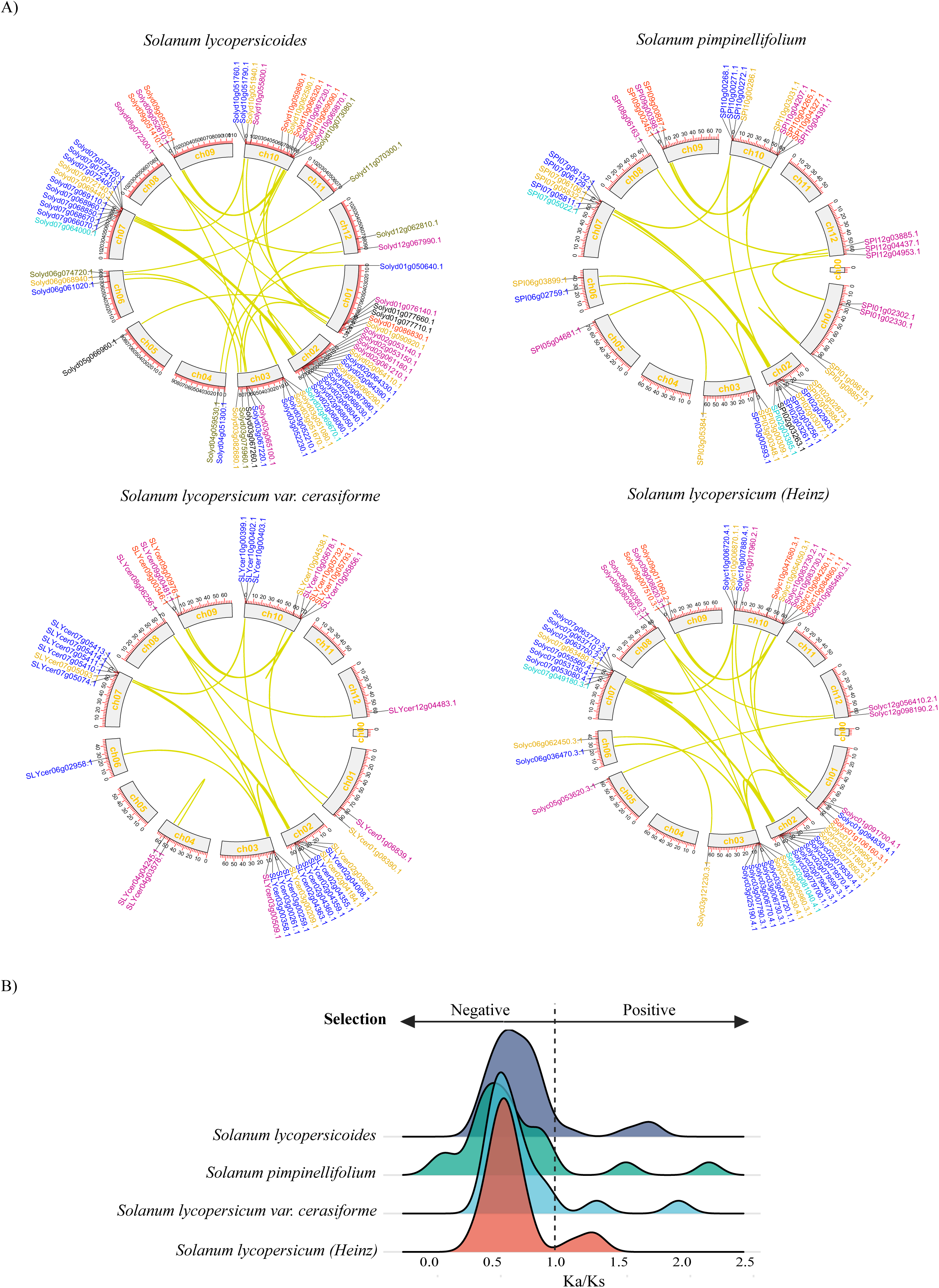
Intra-genomic synteny of lectin genes in *Solanum lycopersicum* (Heinz) and related *Solanum* species. Circos plots illustrate the syntenic relationships of lectin gene family members within the genomes of *S. lycopersicum* (Heinz) and its relatives. Syntenic gene pairs were identified using MCSanX and are shown linked across corresponding chromosomal positions. Lectin genes are color-coded according to their respective gene families, consistent with the scheme used in Figure 1. The size of each chromosome (in megabases) is represented proportionally by the length of its corresponding arc in the plot.

#### Cross-species comparative genomic analysis

To gain deeper insights into the evolutionary history of lectin genes in tomato, we compared gene counts across four *Solanum* genomes—representing both wild and cultivated species—with two ancestral plant genomes, *Amborella trichopoda* and *Vitis vinifera*. This comparison revealed several lineage-specific expansions and contractions in *Solanum*, some of which likely stem from ancient whole-genome duplication (WGD) events, particularly those occurring around the core eudicot WGT (γ event) (Figure S3). In contrast, the relatively stable lectin gene numbers in the ancestral genomes indicate that the diversification seen in tomato is largely a result of more recent duplication and selective processes. Interestingly, cultivated tomatoes displayed few additional expansions relative to their wild relatives, possibly reflecting adaptations linked to domestication. Domain architecture analysis of lectin genes in *A. trichopoda*, *V. vinifera*, and *S. lycopersicum* (cv. Heinz) revealed the emergence of novel functional domains in combination with tomato lectins—including cytochrome P450 (GNA family), NTP_bind (Legume), TIR (Nictaba) and RX_CC (JRL), suggesting functional diversification in these expanded gene families (Figure S4-5; Table S2). Several of these domains are associated with plant defense, such as TIR-Nictaba conferring resistance to spider mites (Santamaria et al., 2019) and RX_CC domains linked to virus resistance (Rairdan et al., 2008). However, to confirm these patterns and distinguish evolutionary events from annotation biases, we further conducted detailed comprehensive orthology and gene family evolution modeling.

Collinearity analysis between *V. vinifera* and *Solanum* genomes identified 772 orthologous lectin gene pairs, revealing extensive retention of multi-copy lectin genes in wild species like *S. lycopersicoides* and *S. pimpinellifolium*. In contrast, the cultivated tomatoes (*S. lycopersicum var. cerasiforme* and *S. lycopersicum* cv. Heinz) showed reduced multi-copy retention, likely reflecting domestication-associated gene loss (Figure 4A, left). Orthogroup clustering using OrthoFinder identified 298 orthogroups containing lectin genes, of which 158 were conserved between *V. vinifera* and *Solanum*, suggesting shared ancestral origins and core functions. The remaining 140 orthogroups were specific to *Solanum*, likely representing lineage-specific duplications or neofunctionalized paralogs that emerged post-divergence and may contribute to species-specific adaptations (Figure 4A, right; Table S10). Phylogenetic reconstruction of lectin genes across *V. vinifera*, wild, and cultivated tomatoes revealed distinct clades comprising both conserved and *Solanum*-specific orthogroups (Figure 4B). Notably, all major lectin families were represented in both conserved and *Solanum*-specific orthogroups, but the JRL and Legume lectin families were disproportionately enriched in the latter.

**Figure 4.**
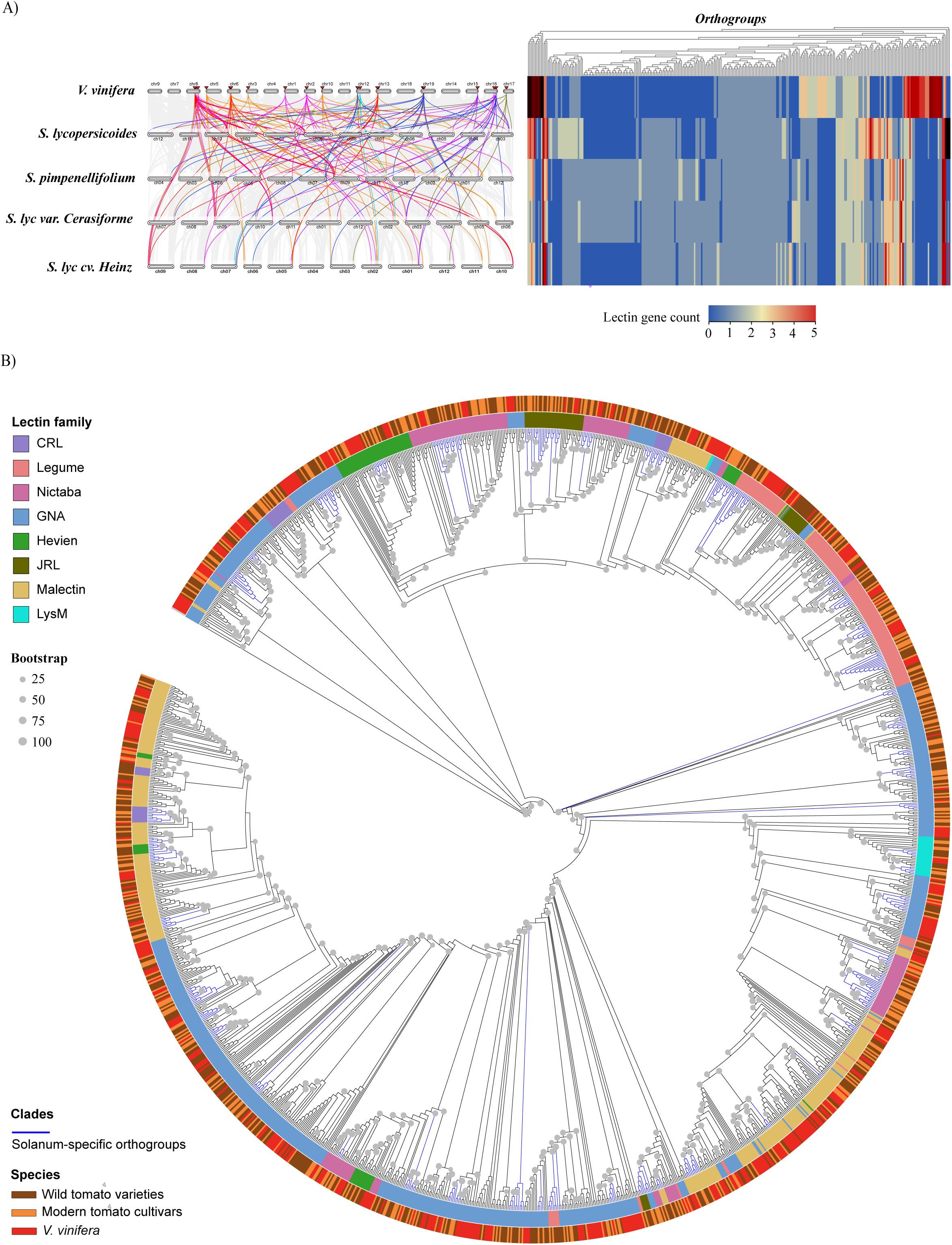
Comparative genomic analysis of the lectin gene family in *Vitis vinifera* and *Solanum species*. A) Intergenomic synteny and orthogroup clustering of lectin genes between *V. vinifera* and four *Solanum* genomes. The left panel shows an intergenomic synteny map with collinear lectin genes linked by color-coded lines, indicating conserved genomic regions across species. The right panel presents a heatmap of orthogroup clusters, highlighting variation in lectin gene counts per orthogroup across the species. Color intensity reflects the number of genes within each orthogroup. B) Rooted maximum likelihood phylogenetic tree illustrating both ancestral conservation and lineage-specific diversification of lectin genes. The first outer color strip denotes gene family classification, while the second strip indicates species origin. Colored downward-pointing triangles mark orthologous gene pairs between *V. vinifera* and *S. lycopersicum* (Heinz) within identified orthogroups. Bootstrap support is represented by the size of black dots at the nodes.

To explore the extent of gene family expansion and contraction, we modeled gene family evolution using a birth-death model of gene gain/loss along the species tree. The analysis identified significant (P < 0.05) expansions and contractions in lectin orthogroups across *Solanum* species using *V. vinifera* as the reference (Figure 5A; Table S11). *S. lycopersicoides* exhibited the highest number of expanded orthogroups (25) and 13 contractions, reflecting retention of duplicated genes. Conversely, *S. pimpinellifolium* and *S. lycopersicum var. cerasiforme* showed fewer expansions (5 and 3, respectively) but a higher number of contractions (10 and 14), suggesting gene loss events during recent evolutionary transitions. Interestingly, *S. lycopersicum* (Heinz) showed a relatively balanced pattern with 10 expanded and 9 contracted orthogroups. We selected the significantly expanded orthogroups in Heinz for in-depth analysis of domain architecture and selective pressure (Figure 5B–C; Table S12). Tomato genes from the GNA, Legume, Hevein, CRL, and Nictaba lectin families that underwent expansion frequently exhibited domain shuffling or gain in their protein architectures. Multiple genes acquired additional domains not present in their ancestral orthologs, such as NTP-binding domain in Legume lectin (*Solyc09g011060*), TIR domains in Nictaba lectins (*Solyc12g096900*; *Solyc12g096880*), and EGF-like domains in GNA lectins (*Solyc10g006710*).

**Figure 5.**
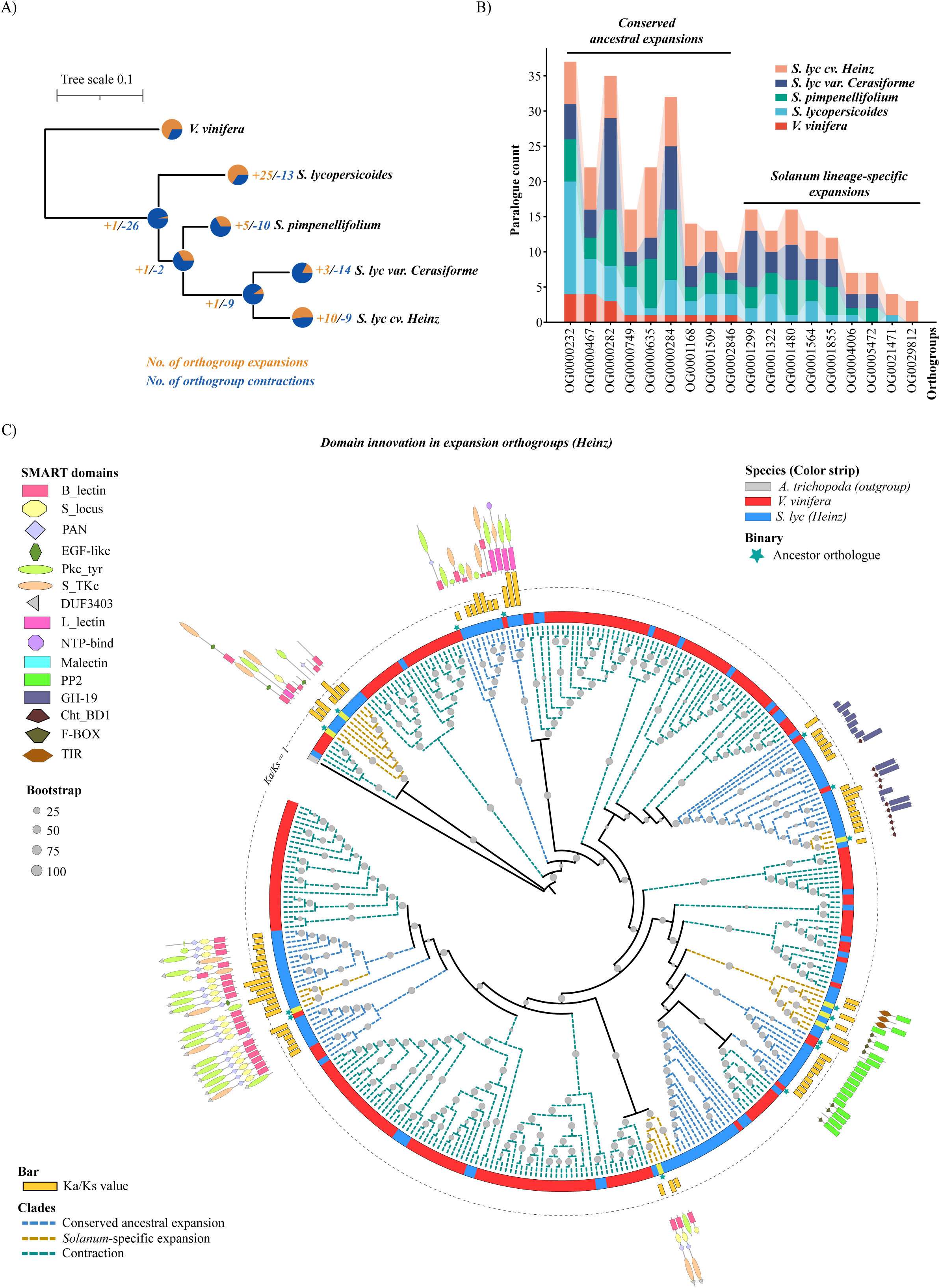
Evolutionary dynamics of orthogroup conservation and protein domain architectures in the lectin gene family. A) Species tree generated using OrthoFinder and CAFE5, depicting expansion (brown) and contraction (blue) of orthogroups containing lectin genes across *Solanum* genomes. The number of expanded or contracted orthogroups is indicated at each node, reflecting evolutionary events. B) Stacked bar plot showing the number of lectin genes in expanded orthogroups, categorized into ancestral expansions (shared with *Vitis vinifera*) and *Solanum*-specific expansions, based on their presence/absence patterns across the genomes. C) Rooted maximum likelihood phylogenetic tree of selected orthogroups illustrating variation in protein domain architectures and patterns of evolutionary selection in *S. lycopersicum* (Heinz) compared to ancestral or wild relative orthologs. The outermost color strip indicates species origin. Branch colors represent ancestral expansions (blue), *Solanum*-specific expansions (brown), and contractions (green). Functional domains are denoted by distinct shapes and colors (legend in top-left corner). Stars mark the ancestral or wild-type orthologs. Yellow bars indicate Ka/Ks values for *S. lycopersicum* genes in expanded orthogroups relative to their ancestral orthologs, with the grey dotted circular reference line denoting a Ka/Ks ratio of 1. Bootstrap support is shown by grey dot sizes at internal nodes.

Although the JRL family was highly represented in *Solanum*-specific orthogroups and showed evident domain addition (e.g., RX_CC gain; Figure S4-5), it did not meet the statistical cutoff for significance in the gene family evolution modeling. This may be due to uneven duplication patterns within the orthogroups or a high baseline number of JRL genes in both ancestral and wild relative genomes, which could obscure detectable rate shifts. Finally, to evaluate whether the expansion of lectin genes across *Solanum* species was accompanied by functional divergence or adaptive evolution, we performed cross-species Ka/Ks analysis. This approach allowed us to compare the evolutionary rates of orthologous genes between *Solanum* species and their ancestral counterparts, helping to identify signatures of selective pressure. The majority of expanded lectin genes exhibited Ka/Ks ratios below 1, indicating purifying selection and suggesting that these genes are under evolutionary constraint to preserve their functions. However, a subset of genes, including those that had acquired domain rearrangements, showed Ka/Ks > 1, pointing toward positive selection. Notably, one Legume lectin gene (*Solyc09g011060*) that gained an NTP-binding domain displayed strong evidence of positive selection, highlighting its potential role in *Solanum*-specific adaptation.

### Expression divergence reveals dynamic evolutionary fates of paralogous lectin genes in cultivated tomato

Previous studies have highlighted that gene duplication in tomato often leads to functional divergence among paralogues (Sato et al., 2012; Kwon et al., 2022; Castaneda et al., 2022). A recent pan-genomics study (Benoit et al., 2025) further emphasized that paralogue retention and diversification in *Solanum* species can have lineage-specific functional implications. GO enrichment analysis of the expanded paralogous genes in tomato revealed significant associations with key biological processes, including MAPK signaling, defense responses, pH regulation, and cell wall organization (Figure S6A). To explore how these duplicated lectin genes have evolved in *Solanum lycopersicum* cv. Heinz, we analyzed their expression dynamics across different developmental tissues. Specifically, we focused on paralogue pairs from expanded orthogroups and examined both their tissue-specific expression levels and co-expression correlations. We also validated the expression and co-expression of homologous lectin pairs in four phylogenetically distant *Solanum* species (*S. macrocarpon*, *S. candidum*, *S. quitoense*, and *S. prinophyllum*) that possess high-quality transcriptomic datasets (Benoit et al., 2025) (Figure 6A–C; Table S13–S17).

**Figure 6.**
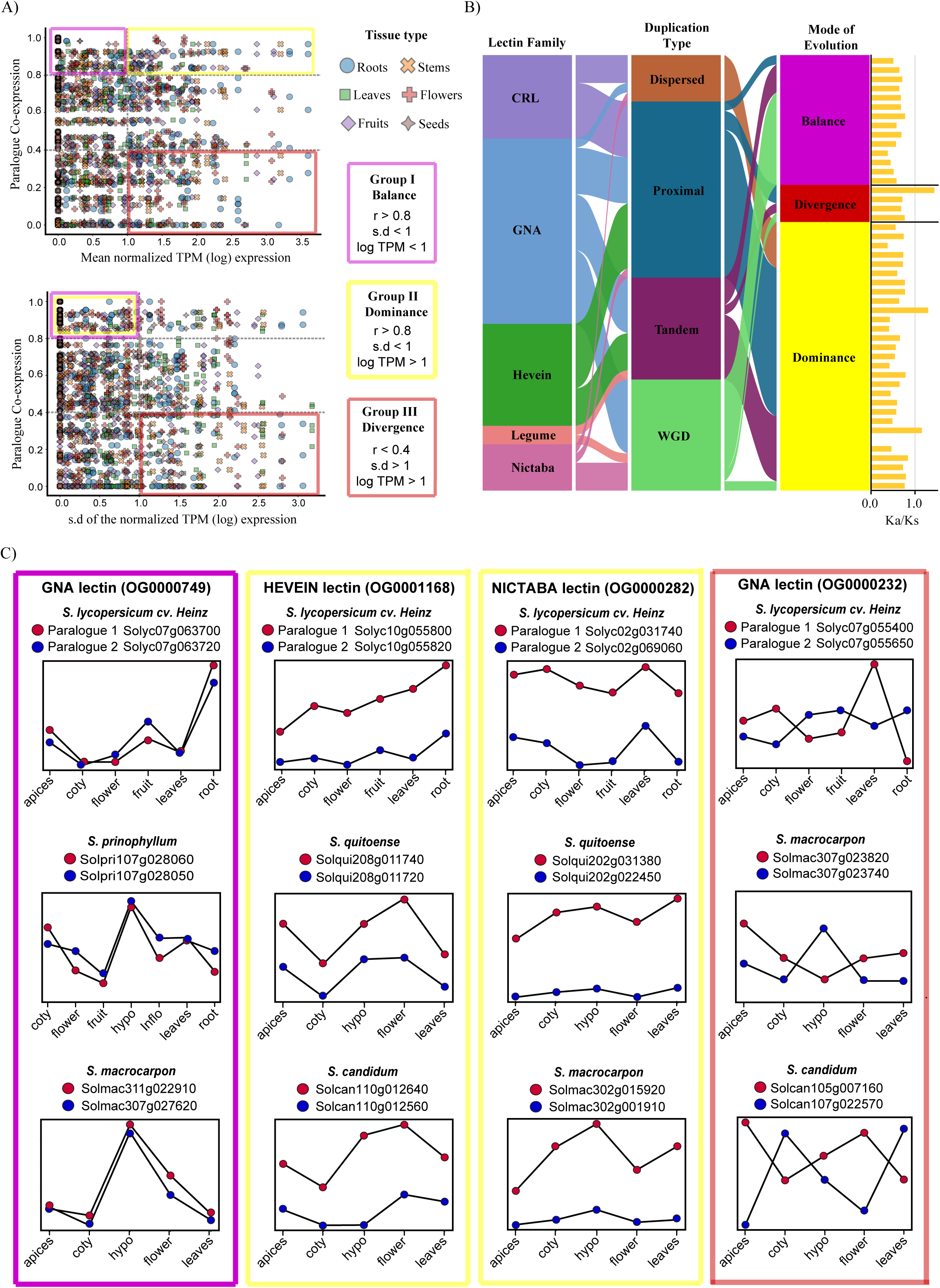
Expression-based functional diversification of paralogous lectin genes in expanded orthogroups. A) Scatter plots showing the distribution of retained paralogous gene pairs in *S. lycopersicum* (Heinz) based on co-expression levels and expression variability. The left panel plots paralogue co-expression (y-axis) against the mean log-transformed normalized TPM (x-axis), while the right panel plots co-expression against the standard deviation (σ) of log-transformed normalized TPM across multiple tissues. Paralogue pairs are categorized into three functional expression groups: balance (pink), dominance (yellow), and divergence (red), based on thresholds of co-expression (r), mean expression, and variability. These categories are summarized in the schematic at the top. B) Alluvial plot depicting the interplay between duplication type, expression-based evolutionary modes, and selective pressure (Ka/Ks ratio) among paralogous gene pairs classified under individual lectin families. C) Representative examples of *S. lycopersicum* paralogue pairs illustrating the three distinct expression divergence patterns, also conserved in other *Solanum* relatives (*S. macrocarpon*, *S. candidum*, *S. quitoense*, and *S. prinophyllum*). Colored boxes indicate the functional category (as defined in panel A). Red and blue circles denote the expression levels of paralogue 1 and paralogue 2, respectively, across various developmental tissues, including apices (shoot apices), cotyledons (coty), hypocotyls (hypo), inflorescence (inflo), flower, fruits, and roots.

Based on expression correlation and variance across tissues (Figure 6A), we classified the paralogue pairs into four major evolutionary expression categories (Figure 6B-C): (i) *Balance* – where both genes are expressed at similar levels across tissues, suggesting conservation of total gene dosage and function; (ii) *Dominance* – where one paralogue consistently shows higher expression than its counterpart, indicating asymmetric contribution possibly reflecting early signs of subfunctionalization or pseudogenization; (iii) *Specialization* – where paralogues exhibit tissue-specific expression, implying potential neofunctionalization or subfunctional roles; and (iv) *Divergence* – where expression levels and patterns differ substantially across all tissues, suggesting complete functional divergence post-duplication.

In total, we identified 14 paralogue pairs in the *balance* group, 29 in *dominance*, 3 in *specialization*, and 4 in *divergence*. However, the expression patterns of Heinz paralogue pairs in the *specialization* group were not consistent with their homolog expression in distant relative solanum species. In addition, A large number of paralogue pairs (118) could not be confidently classified into any group due to ambiguous or inconsistent expression patterns across tissues and species. These were designated as an *Undefined group* (Table S13). This undefined category may reflect technical limitations such as incomplete transcript coverage, low or context-specific expression, or biological factors such as conditional or epigenetic regulation. Future investigations incorporating broader developmental stages, stress responses, or single-cell resolution may help resolve their evolutionary trajectories.

Interestingly, most paralogue pairs in the defined groups originated from conserved orthogroups, suggesting evolutionary constraints that maintain gene function across species (Table S13). In contrast, nearly all paralogues from *Solanum*-specific orthogroups fell into the undefined group, possibly reflecting recent duplication events with unresolved or highly dynamic regulatory evolution.

We further investigated the relationship between duplication mechanisms and expression divergence. Notably, the majority of paralogue pairs in the balance group were derived from WGD, consistent with retention of gene dosage and function typically associated with whole-genome or large-scale events. In contrast, paralogue pairs in the dominance and divergence groups were primarily associated with small-scale duplications (SSDs) such as tandem, proximal, and dispersed duplication (Figure 6B; Table S13). These types of local duplications often generate new gene copies that are less constrained by ancestral dosage balance and therefore more prone to regulatory divergence. Interestingly, most SSD-derived paralogues were also part of conserved orthogroups, indicating that even after speciation, ancestral lectin genes have undergone recent local duplications, with some new copies being selectively retained and showing signs of functional divergence (Table S13).

Consistent with this interpretation, Ka/Ks analysis revealed that paralogues in the balance group were under strong purifying selection (Ka/Ks < 1), maintaining sequence conservation, while some paralogues in the divergence and dominance groups showed Ka/Ks > 1, indicating adaptive evolution (Figure 6B; Table S13). For instance, a GNA family pair (*Solyc07g055400:Solyc07g055650*) in the divergence group had a Ka/Ks of 1.43 (p = 0.00003), and a Hevein family pair (*Solyc10g074440:Solyc10g074460*) in the dominance group had a Ka/Ks of 1.15 (p = 0.0006). These results suggest that positive selection, in combination with small-scale duplication, may be driving functional divergence in specific lectin gene families.

Collectively, this analysis underscores the dynamic evolutionary paths of lectin gene duplicates in tomato. WGD appears to maintain essential functions through dosage conservation, whereas small-scale duplications contribute to functional divergence, even within gene families in conserved orthogroups. The high proportion of dominant and undefined expression patterns points to an ongoing process of regulatory fine-tuning and evolutionary experimentation. Moreover, the convergence of duplication type, expression divergence, and Ka/Ks patterns strengthens the case for local duplications as a key source of functional innovation post-speciation.

### VIGS-mediated silencing of *Solyc04g077390* and *Solyc07g063700* reduces disease progression of *Ralstonia solanacearum* in *Solanum lycopersicum (Heinz)*

For functional validation, we selected two *GNA* lectin genes: one singleton (*Solyc04g077390*) and one whole-genome duplication (WGD)-derived paralogue (*Solyc07g063700*). The latter belongs to a balanced group (Figure 6B) that exhibits conserved developmental expression across tomato and four phylogenetically distant relatives, suggesting preservation due to essential ancestral function. Notably, both genes were induced upon *Ralstonia solanacearum* infection (Figure S6B-C). To assess their role, we employed a tobacco rattle virus (TRV)-based virus-induced gene silencing (VIGS) approach in *Solanum lycopersicum*. The efficiency of VIGS was confirmed using *Phytoene Desaturase* (PDS) as a positive control, which exhibited characteristic photobleaching symptoms 17 days post-infiltration (Figure S7B). Quantitative analysis confirmed substantial downregulation of the target genes, with *Solyc04g077390* and *Solyc07g063700* showing ∼90% and ∼70% reduction in transcript levels, respectively (Figure S7C).

Following successful gene silencing, *R. solanacearum* was inoculated (ODCCC = 0.1) via the soil drench method into both the VIGS vector control (pTRV:00) and knockdown plants (Chandan et al., 2023). Disease severity was evaluated using a wilting index over seven days. By the seventh day post-infection, wilting in vector control plants reached 98.32%, whereas knockdown lines of *Solyc04g077390* and *Solyc07g063700* exhibited significantly reduced wilting at 64.73% and 67.42%, respectively. Statistical analysis using the Friedman and Duncan tests confirmed the significance of these observations, and the Wilcoxon signed-rank test further supported that disease progression was significantly lower in the knockdown lines (Figure 7A– B). To quantify bacterial proliferation, colony-forming units (CFUs) were measured in both roots and shoots. While no significant differences in root CFUs were observed between control and silenced plants, shoot tissues of the knockdown lines harbored significantly fewer bacteria compared to the vector control (Figure 7C). These results suggest that silencing of *Solyc04g077390* and *Solyc07g063700* impairs systemic colonization of *R. solanacearum*, particularly in aerial tissues. Given that *R. solanacearum* primarily colonizes xylem vessels (Caldwell et al., 2017), we examined vascular changes in infected crown stem sections. Vector control plants exhibited a ∼65.45% increase in xylem area compared to uninfected controls. In contrast, knockdown lines of *Solyc04g077390* and *Solyc07g063700* showed markedly reduced xylem expansion of ∼22.1% and ∼3.8%, respectively (Figure 7D; Figure S7D), further supporting attenuated infection and spread in these lines.

**Figure 7.**
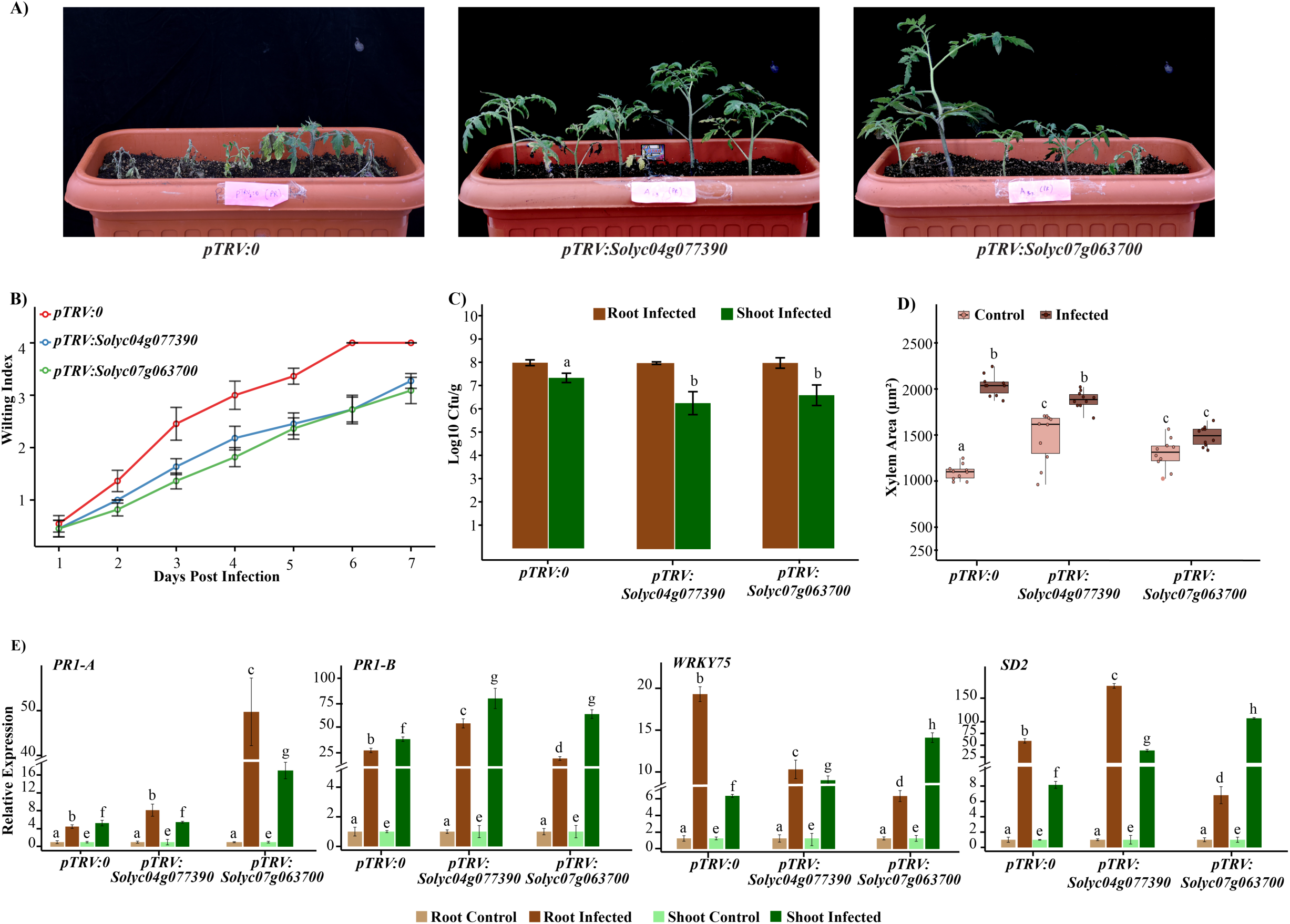
Silencing of *Solyc04g077390* and *Solyc07g063700* enhances resistance to bacterial wilt in *Solanum lycopersicum*. A) Phenotypic comparison of VIGS (virus-induced gene silencing) plants targeting *Solyc04g077390* and *Solyc07g063700* following infection with *Ralstonia solanacearum*. B) Line plot showing the wilting index over time in infected plants, indicating delayed disease progression in silenced lines compared to control. C) Quantification of bacterial load (colony-forming units) in stem tissue at 7 days post inoculation, showing reduced pathogen proliferation in silenced plants. D) Boxplot representing the increase in xylem bundle area in infected plants. The central line indicates the median, while box limits denote the first and third quartiles. E) Relative expression levels of defense-related genes (*PR1A*, *PR1B*, *WRKY75*, and *SD2*) in control (pTRV:00) and silenced lines (pTRV:*Solyc04g077390* and pTRV:*Solyc07g063700*). Different letters denote statistically significant differences between groups (p < 0.05).

To elucidate the downstream defense responses modulated by these genes, we analyzed expression patterns of key defense-related genes, including *SlPR1-a*, *SlPR1-b*, *SlWRKY75*, and *SlSD2* (Figure 7E). These genes are known to participate in salicylic acid- and ethylene-mediated defense signaling and are activated upon *R. solanacearum* infection (Hamilton et al., 2024; Chen et al., 2022; Yang et al., 2024). In the *Solyc07g063700*-silenced line, *SlPR1-a* showed strong upregulation, with a 49.72-fold increase in the root and 16.99-fold in the shoot. Although root transcript levels of *SlPR1-b* were inconclusive, its expression in the shoot was markedly elevated: an 80.71-fold increase in the *Solyc04g077390*-silenced line and a 63.9-fold increase in the *Solyc07g063700*-silenced line, compared to a 37.00-fold increase in the vector control. Interestingly, *SlWRKY75* transcript levels in roots were higher in the vector control (19.03-fold) than in the *Solyc04g077390* (10.04-fold) and *Solyc07g063700* (6.04-fold) knockdown lines. However, in shoot tissues, *SlWRKY75* expression was significantly higher in silenced plants (8.75-fold and 13.83-fold for *Solyc04g077390* and *Solyc07g063700*, respectively) compared to the control (6.08-fold). Expression of *SlSD2*, a defensin gene, in root tissues showed no consistent trend. In contrast, shoot tissues exhibited significantly higher *SlSD2* expression in knockdown lines: 38.9-fold in *Solyc04g077390* and 107.2-fold in *Solyc07g063700*, compared to only an 8.13-fold increase in the vector control (Figure 7D).

Together, these findings indicate that silencing of *Solyc04g077390* and *Solyc07g063700* enhances resistance to *R. solanacearum* by limiting bacterial spread, suppressing xylem colonization, and modulating the expression of key defense-related genes. These genes likely act as negative regulators of defense and represent promising targets for improving disease resistance in tomato.

## Discussion

Plant genome evolution operates on two interconnected levels: the evolution of individual gene families and that of entire genomes (Clark, 2023). Over evolutionary time, gene families have undergone expansions and contractions, contributing to both structural and functional conservation or diversification—particularly among genes involved in critical developmental processes (Kodama et al., 2022; Streubel et al., 2023). Lectins, known for their carbohydrate-binding properties, exemplify this structural diversity. Their sugar and ligand-binding capabilities have been shown to play essential roles in plant development (Peng et al., 2020; Petrova et al., 2021) and in responses to various biotic and abiotic stresses (Stefanowicz et al., 2016; Pan et al., 2020; Yuan et al., 2025). It is proposed that lectin functions diversified throughout plant evolution at both the nucleotide and protein levels (Van Holle et al., 2017; Van Holle and Van Damme, 2019). In this study, we selected tomato (*Solanum lycopersicum*) as our model system due to its agricultural relevance and the availability of comprehensive genomic resources. Our goal was to investigate the evolutionary and functional dynamics of lectin gene families in the tomato genome.

### Structural diversification and functional relevance of multi-domain tomato lectins

Focusing first on domain occurrence, we analyzed 12 lectin domain-containing families, most of which are not unique to the plant kingdom but are also found in other evolutionary lineages such as bacteria and fungi. The notable exceptions are the Amaranthin and *Euonymus europaeus* agglutinin (EUL) domains, which are restricted to plants (Van Holle and Van Damme, 2019). In the tomato genome, we identified eight lectin families; the Amaranthin, EUL, Ricin-B, and Agaricus bisporus agglutinin (ABA) families were absent. The ABA family is thought to have originated in fungi and is largely confined to fungal species, with limited representation in certain bryophytes (Van Holle and Van Damme, 2019). Although Amaranthin and EUL are bona fide plant lectins, they represent relatively small gene families. The Amaranthin domain, in particular, shows taxonomic restriction—being enriched in Poaceae and Rosales species, but nearly absent in the Asterid clade, to which tomato belongs (Dang et al., 2017). This suggests that some lectin domains may be lineage-specific and potentially adapted to perform specialized roles in development or stress responses (Domazet-Lošo et al., 2024).

Our findings revealed considerable structural diversity among lectin gene families in tomato. Many of these lectin genes encode multi-domain proteins, in which lectin domains are integrated with various other annotated functional domains. Previous studies have indicated that most lectin domain architectures originated in early ancestors of *Streptophyta* or *Chlorophyta* and have been conserved throughout evolution (Van Damme et al., 2017). These multi-domain arrangements are likely not solely structural but may provide adaptive functional advantages. For example, chimeric proteins containing GNA domains have been shown to enhance resistance against rice blast and *Fusarium* wilt in tomato (Chen et al., 2006; Catanzariti et al., 2015). Interestingly, a T-DNA insertion mutant of a GNA chimerolectin in *Arabidopsis* was reported to act as a negative regulator of plant immunity during root-knot nematode infection, with its downregulation leading to reduced gall formation (Zhou et al., 2023). In our study, among more than 50 identified GNA chimerolectins, a subset—including *Solyc04g077320*— was induced, while others such as *Solyc03g006730*, *Solyc10g005440*, *Solyc01g014560*, and *Solyc02g079690* were downregulated upon root-knot nematode infection, despite sharing similar domain architectures. Notably, *Solyc02g079690* and *Solyc04g077320* also exhibited induction in the root meristem. Since root-knot nematodes typically penetrate host plants via the root tip, the differential regulation of *Solyc04g077320* and *Solyc02g079690* in root meristem tissues suggests their potential involvement in modulating localized immune responses at the primary site of nematode invasion.

The widespread abundance of chimerolectins across lectin families can largely be attributed to their versatile roles in pathogen defense and signaling. However, we observed notable exceptions within the JRL and Hevein lectin families, where hololectins were more prevalent. The functional advantage of multiple repeated lectin domains within hololectins remains incompletely understood. Nevertheless, functional evidence from rice supports the potential significance of JRL hololectins. For instance, overexpression of the JRL hololectin gene *Orysata* (also known as *SALT*), located within the major quantitative trait locus (QTL) Saltol-1, has been shown to confer multiple stress responses, including enhanced salinity tolerance, negative regulation of cold tolerance, and increased resistance against *Magnaporthe grisea* (Shinjo et al., 2011; He et al., 2017; An et al., 2024). In our analysis, the JRL hololectins *Solyc04g080410* and *Solyc04g080420* each contained two jacalin-like domains, while *Solyc10g078600* harbored three jacalin-like domains. Notably, *Solyc10g078600* exhibited complex regulation across multiple stresses: it was downregulated under endoplasmic reticulum stress, induced under cold stress, and differentially regulated during salt stress progression. Whether these multiple jacalin-like domains function synergistically or conditionally in mediating abiotic stress responses remains to be functionally elucidated.

Similarly, chitin-binding Hevein hololectins are abundant in cereals and have been extensively studied for their agglutination activity (Mishra et al., 2019). Representative examples include wheat germ agglutinin (WGA) and *Oryza sativa* agglutinin (OSA). Interestingly, WGA, which contains multiple Hevein domains, not only exhibits strong affinity for fungal chitin but also binds sialic acid, contributing to its implication in celiac disease (De Punder et al., 2013). This illustrates that even structurally similar lectin domains may exhibit distinct binding specificities under different physiological conditions. In tomato, while Hevein chimerolectins were expressed under fungal stress conditions, the four identified Hevein hololectins (*Solyc03g116190*, *Solyc11g020130*, *Solyc11g020450*, *Solyc11g020530*) did not exhibit significant expression in available stress or developmental transcriptomic datasets. This absence of detectable expression may reflect transient or tightly regulated induction, possible deregulation, or a shift in functional relevance following monocot-dicot divergence, where such proteins may have retained greater functional significance in monocots (Van Damme et al., 2017). Overall, whether the presence of multiple repeated domains confers functional advantages remains poorly characterized, and direct functional comparisons between hololectins, merolectins, and chimerolectins are still lacking.

Furthermore, our comparative analysis between the tomato genome and ancestral genomes uncovered several domain combinations not previously observed in basal or distant angiosperm ancestors. Some tomato lectin genes appear to have acquired additional domains such as TIR, NTP-bind, and RX_CC, leading to the formation of new chimerolectins likely to contribute to functional diversification. For example, an Arabidopsis Nictaba-TIR chimera has been implicated in enhanced resistance to spider mites via transcriptional reprogramming (Santamaria et al., 2019). Although RX_CC domains are individually well known for conferring resistance to pathogens such as *Potato virus X* (Rairdan et al., 2008), and NTP-binding domains are implicated in the intercellular movement of RNA viruses (Wu and Cheng, 2020), there is currently no direct functional evidence supporting the roles of chimeric proteins containing JRL-RX_CC or Legume-NTP_bind domain architectures in stress tolerance. Notably, our meta-transcriptomic analysis revealed specific induction of *Solyc09g090445* (JRL-RX_CC) during anthracnose disease progression suggesting a role in mediating responses to fungal infections. However, no significant expression was observed for *Solyc09g011060* (Legume-NTP_bind) under any stress conditions in our meta-transcriptomic datasets.

Collectively, the structural and meta-transcriptomic analyses have identified previously uncharacterized chimerolectin candidates in tomato that exhibit distinct expression patterns in response to specific diseases, including root-knot nematode and anthracnose, as well as in developmental tissues that may serve as key entry points for pathogens and facilitate disease progression. Moreover, several uncharacterized hololectin candidates were identified, providing opportunities to investigate the potential functional advantages or disadvantages associated with the presence of repeated lectin domains under various stress conditions.

### Functional diversification of lectin paralogues during tomato evolution

All dicotyledonous plants share the first WGT event (γ event) that has been linked to diversification of eudicots and angiosperms (De Bodt et al., 2005). In addition to the γ WGT event, Solanaceae species experienced a lineage-specific WGT event (*T* event) approximately 65 million years ago (Sato et al. 2012). Although the duplicated genes have since diverged, three distinct tomato sub-genomes remain identifiable when compared to species such as *Vitis vinifera*, which only underwent the γ event (Huang et al., 2022). These sequential polyploidy events are likely to have shaped the structure and function of lectin genes and their paralogues in the tomato genome. Based on our collinearity, orthology, and phylogenetic analysis, the impact of these events on lectin gene family expansion can be conceptualized in two major ways: (i) *Conserved ancient expansion* - lectin paralogues that originated from the γ event and possess ancestral orthologs may have been either retained in their original form or modified by the subsequent *T* event; and (ii) *Lineage-specific expansion* - the *T* event may have led to the emergence of additional lectin paralogues that lack identifiable ancestral orthologs.

Comparative analysis of the *Vitis vinifera* and *Solanum* genomes reveals that both WGT events were followed by extensive gene loss within the lectin family—an outcome consistent with the action of purifying selection, which acts to eliminate redundant or deleterious duplicates (Ohno, 1970). Nevertheless, our findings indicate notable retention and expansion of specific lectin families in tomato—particularly GNA, Nictaba, Legume, CRL, and Hevein— predominantly under purifying selection, with a few members showing signatures of positive selection. Interestingly, both conserved and lineage-specific lectin paralogues exhibit substantial variation in domain architecture, including domain gains, losses, and rearrangements. These structural innovations are likely driven by a combination of whole-genome duplications and small-scale duplication mechanisms, such as tandem, proximal, and dispersed duplications. According to the dosage balance hypothesis, genes in multiprotein complexes must retain stoichiometric balance for optimal function. Whole-genome duplications (WGDs) preserve this balance by duplicating all genes simultaneously, aiding duplicate retention (Birchler and Veitia, 2014; Conant et al., 2014). In contrast, small-scale duplications (SSDs) can disrupt dosage balance, often gradually leading to gene inactivation or loss (Panchy et al., 2016). However, regardless of its type, duplicated genes with beneficially increased expression during different developmental or environmental responses may be retained longer (Van de Peer et al., 2017). In flowering plants including Solanaceae genomes, it has been observed that many gene duplicates tend to return to single-copy status following WGD events. However, gene families involved in stress responses, signaling, and development often retain multi-copy status, likely due to their functional relevance under specific physiological or environmental conditions (Maere et al., 2005; Li et al., 2016; Zhang et al., 2025). For instance, in our analysis, we found that tandem duplicates from the Hevein lectin family (*Solyc10g055800*; *Solyc10g055810*; *Solyc10g055820*) belonging to the conserved ancient expansion group showed highly specific induction during anthracnose disease progression and fruit development, highlighting a possible retention due to its role in fruit-specific defense response. In this context, our study raises an important question: Do the expanded lectin paralogue pairs resulting from WGD or SSD exhibit functional biases based on their expression in stress conditions or in specific developmental tissues?

To investigate the functional dynamics of lectin gene duplicates, we analyzed their expression and co-expression profiles across various developmental tissues in tomato and four phylogenetically distant relatives. This comparative approach enabled us to evaluate how different modes of gene duplication influence the evolutionary trajectories of paralogous expression (Benoit et al., 2025). Paralogue lectin pairs originating from small-scale duplications (SSDs), particularly tandem and proximal events, frequently exhibited divergent expression patterns—either through spatial expression differences or through one paralogue displaying dominant expression throughout development in both tomato and its relatives. In contrast, paralogues derived from whole-genome duplications (WGDs) generally retained similar expression profiles across all developmental stages, indicating greater functional conservation. Our interpretations are consistent with findings in *Arabidopsis*, where the extent of functional divergence among gene duplicates was shown to be influenced by the duplication mechanism. Ezoe et al. (2021) reported that tandemly duplicated genes tend to diversify functionally, whereas WGD-derived genes often retain ancestral functions. This was further supported by single and double knockout mutant analyses, where morphological changes confirmed functional specialization among certain paralogues.

Although the lack of stress-responsive transcriptomic datasets in distant relatives limited parallel expression analyses under stress conditions, Gene Ontology (GO) enrichment of expanded lectin paralogues in tomato revealed enrichment for key biological processes such as MAPK signaling, pH regulation, cell recognition, and defense responses. This supports the hypothesis that paralogues, especially those retained from SSDs, may be retained due to their involvement in adaptive responses and signaling pathways.

### Functional validation of GNA chimerolectins

To investigate the functional relevance of lectin genes, we selected two GNA chimerolectins — one singleton and one WGD-derived paralog from the balanced group—for functional validation. GNA chimerolectins have previously been implicated in defense responses against bacterial and fungal pathogens in tobacco and strawberry (Kim et al., 2015; Ma et al., 2022). The two selected genes, *Solyc04g077390* and *Solyc07g063700*, encode proteins containing a B-type lectin domain, an S_locus_glycoprotein domain, and a PAN domain. The B-type lectin domain facilitates mannose binding during bacterial infection, while the S_locus_glycoprotein and PAN domains are associated with immune modulation (Maimbo et al., 2010; Kim et al., 2015). Notably, the conserved PAN domain has been shown to suppress immune signaling by downregulating jasmonic acid, ethylene, and MAPK pathways, while promoting the expression of negative regulators such as *NINJA*, *HDA6*, and *TOPLESS*. Conversely, mutations in this domain trigger enhanced immune responses and inhibit *Botrytis cinerea* invasion (De et al., 2023).

Using Virus-Induced Gene Silencing (VIGS), we found that silencing *Solyc04g077390* and *Solyc07g063700* improved plant performance compared to control plants harboring the empty vector. Knockdown lines exhibited significantly increased expression of defense-related genes—*PR1A* (in *Solyc07g063700*) and *PR1B*, *WRKY75*, and *SD2* (in both lines)—as well as a reduced bacterial load, reflected by lower CFU counts in shoot tissues. This enhanced immune response was also associated with fewer wilting symptoms. Interestingly, control plants showed a larger xylem area, likely as a compensatory response to xylem blockage caused by higher bacterial colonization (Ingel et al., 2022). In contrast, knockdown lines had less xylem expansion, consistent with reduced vascular stress due to lower pathogen load.

These findings suggest that the two GNA chimerolectins contribute to susceptibility during bacterial wilt attack. When comparing these chimerolectins to GNA merolectins—lectins shown to confer resistance—we observed key differences in domain architecture and sequence composition. Specifically, the chimerolectins uniquely possessed an S_locus_glycoprotein domain and showed sequence variations within the conserved PAN domain, which were absent in the resistance-associated merolectins. These structural changes may play a pivotal role in modulating immune responses. Further investigation into how the acquisition of additional functional domains alters lectin function could provide important insights into the evolution of plant immunity.

## Conclusion

This study provides an integrative view of the structural, evolutionary, and functional landscape of lectin gene families in tomato. The remarkable diversity in domain architectures, particularly among chimerolectins and hololectins, underscores their evolutionary plasticity and potential for functional innovation. Our comparative genomic and expression analyses reveal that whole-genome and small-scale duplications have jointly shaped lectin gene family expansion, with distinct expression divergence patterns reflecting evolutionary trajectories. Functional assays confirmed that specific GNA chimerolectins act as negative regulators of defense, likely through domains that suppress immune signaling. Our findings advance our understanding of lectin evolution, functional diversification and identify promising candidates for engineering disease resistance in tomato and other Solanaceae crops.

## Author contributions

**Vishnu Shukla:** Conceptualization, data curation, formal analysis, investigation, visualization, writing—original draft, writing—review and editing. **Abin Panackal George:** formal analysis, Conducting VIGS experiments, investigation, visualization, writing—original draft, writing—review and editing. **Rama Sai Venkata Marthi:** formal analysis, VIGS investigation, RT-qPCR validations, writing—review and editing. **Abhilasha Prashant Sonawane:** Meta transcriptome expression analysis, writing—review and editing. **Sanskruti Parida:** stress experiments, RT-qPCR validations, writing—review and editing. **Eswarayya Ramireddy:** Conceptualization, data curation, investigation, visualization, methodology, funding acquisition, project administration, writing—original draft, writing—review and editing.

## Funding

This work was supported by IISER Tirupati and by an Early Career Research Award from the Science and Engineering Research Board, Department of Science and Technology, Government of India (ECR/2016/001071) awarded to E.R. Additional support was provided by the Ignite Grant [AgSci/24-25/03] from the Ignite Life Science Foundation to E.R. V.S. and A.P.G. acknowledge funding from IISER Tirupati for postdoctoral and graduate studies, respectively. R.S.V.M. acknowledges fellowship support from the Ignite Grant, and S.P. acknowledges the CSIR-JRF fellowship.

## Acknowledgments

E.R. gratefully acknowledges the support of IISER Tirupati, particularly the central facilities. We thank Prof. Suvendra Kumar Ray (Tezpur University) for providing the *Ralstonia solanacearum* F1C1 strain. Dr. Gopaljee Jha (NIPGR), for helping the VIGS experiments.

## Data availability statement

All relevant experimental data can be found within the manuscript and its supporting materials. The Scripts and R codes are submitted at GitHUb can be accessed using the following link (https://github.com/ERR-lab/Lectin-gene-family-in-tomato/tree/main/Lectin_manuscript_GitHub_files) .

## Conflict of interest disclosure

The authors declare no conflicts of interest.

## Supplementary Figure legends

**Figure S1: Transcript abundance of lectin genes in *S. lycopersicum* (Heinz) during development.** This heatmap shows the clustering of lectin genes based on their normalized transcript abundance (TPM) across various developmental tissues, including roots, stems, leaves, flowers, fruits, and seeds. Gene family types are color-coded for distinction, and expression levels are represented by the intensity of the color gradient, as indicated in the legend.

**Figure S2: Transcript abundance of lectin genes in *S. lycopersicum* (Heinz) during stress.** This heatmap shows the clustering of lectin genes based on their transcript abundance (log_2_FC) across various biotic and abiotic stresses. Gene family types are color-coded for distinction, and expression levels are represented by the intensity of the color gradient, as indicated in the legend.

**Figure S3. Comparative distribution of lectin gene counts in Solanum species and their ancestral genomes.** This figure presents a comparative map illustrating lectin gene counts across extant *Solanum* genomes and their inferred ancestral lineages. Circle size is proportional to the number of lectin genes in each genome. Color of each circle is based on lectin gene family type. Major evolutionary events are marked with colored downward arrows.

**Figure S4: Protein domain architecture of lectin genes in *S. lycopersicum*, *V. vinifera* and *A. trichopoda***. Based on NCBI-CDD annotations, protein domains were mapped along the lectin sequences. Gene family types are color-coded for distinction. Domain shape and colors for each species is highlighted in the legend.

**Figure S5. Impact of whole-genome triplication and duplication events on protein domain expansions in lectin gene families.** This figure summarizes the expansion of protein domains in *S. lycopersicum* lectin gene families, compared to their counterparts in key ancestral species such as *Amborella trichopoda* and *Vitis vinifera*. It highlights both ancestral and lineage-specific domain acquisitions, underscoring the role of genome duplication events in the structural diversification of lectin genes during evolution.

**Figure S6. Functional roles of expanded lectin genes in *S. lycopersicum* (Heinz) under stress conditions.**

(A) *Left panel*: Bubble plot showing Gene Ontology (GO) enrichment analysis of expanded lectin genes. The x-axis represents the proportion of lectin genes associated with each GO term relative to the total number of genes annotated for that term in the tomato genome. Bubble size indicates the number of lectin genes mapped to each term, while the color gradient denotes statistical significance of enrichment.

*Right panel*: Heatmap illustrating transcript abundance (logC fold change) of expanded lectin genes under various biotic and abiotic stress conditions. Genes are clustered based on expression patterns, with gene family types color-coded and expression intensity represented by a gradient scale.

(A) (B) Relative expression of selected expanded lectin genes and response genes upon *Ralstonia solanacearum* infection. Statistical significance compared to control is indicated (*p ≤ 0.05; **p ≤ 0.01; **p ≤ 0.001; Student’s *t*-test).

**Figure S7. Assessment of virus-induced gene silencing (VIGS) controls for functional validation of selected lectin genes in *S. lycopersicum* during bacterial wilt infection.**

(A) Wilting symptoms observed in the susceptible tomato variety Pusa Ruby at 5 days post infection (DPI), compared to uninfected control plants. This was performed to validate and standardize the infection procedure and symptom development.

(B) Visual phenotypes of VIGS-silenced plants (pTRV:*Solyc04g077390* and pTRV:*Solyc07g063700*) under control (non-infected) conditions, compared to the empty vector control (pTRV:0) and positive control (*pTRV:PDS*).

(C) Bar plot showing the relative reduction in gene expression of *Solyc04g077390* and *Solyc07g063700* in roots and shoots of VIGS-silenced plants compared to pTRV:0 controls. Asterisks indicate statistically significant differences relative to controls (p ≤ 0.001; Student’s *t*-test).

(D) Representative images illustrating increased xylem area in pTRV:*Solyc04g077390* and pTRV:*Solyc07g063700* plants following bacterial wilt infection, compared to pTRV:0 controls.

## Supplementary Table legends

**Table S1: Identification of putative lectin domain containing genes across multiple ancestral and *Solanum* genomes.** HMMER-based domain scan results for *S. lycopersicum* (Heinz), *S. lycopersicoides*, *S. pimpinellifolium*, *S. lycopersicum var. cerasiforme*, *V. vinifera*, and *A. trichopoda*. Gene identifiers are annotated based on the presence of statistically significant hits for various lectin domains, providing a foundational dataset for comparative lectin family analysis.

**Table S2: Protein domain architectural variations in lectin gene families across *A. trichopoda*, *V. vinifera*, and *S. lycopersicum* (Heinz).** The table presents a survey of lectin genes based on NCBI-CDD annotations, highlighting the sequence positions and types of protein domains within different lectin superfamilies.

**Table S3: Gene Ontology (GO) enrichment analysis of lectin genes in Solanum lycopersicum (Heinz).** GO enrichment results obtained using ShinyGO, categorizing lectin genes into enriched biological processes, molecular functions, and cellular components.

**Table S4: Enrichment analysis of transcription factor (TF) binding sites in the promoters of lectin genes in *Solanum lycopersicum* (Heinz).** Promoter regions (2 kb upstream) of each lectin gene were scanned for transcription factor family binding motifs using the AME module of the MEME Suite. Enrichment was assessed by comparing the frequency of TF binding sites in lectin gene promoters to their genome-wide distribution in tomato. Enrichment scores are represented as -logCC (adjusted p-value).

**Table S5: Meta-transcriptomic analysis of lectin gene expression during development in *Solanum lycopersicum* (Heinz).** A total of 22 transcriptomic datasets representing various developmental tissues (roots, stem, leaves, flowers, fruits, and seeds) were analyzed. When normalized TPM values were not available, differential expression analysis was performed using DESeq2. Lectin gene expression was considered significant based on a threshold of log(TPM) > 1 and p < 0.05.

**Table S6: Tissue-specific expression analysis of lectin genes in developmental tissues of *Solanum lycopersicum* (Heinz)**. Developmental expression datasets (Table S5) were used to calculate the tissue specificity index (***τ***) for lectin genes across various developmental tissues, including roots, stems, leaves, flowers, fruits, and seeds. Additionally, analyses were performed for subsets of root-specific, flower-specific, and fruit-specific tissues to assess expression patterns in more specialized contexts.

**Table S7: Meta-transcriptomic analysis of lectin gene expression during biotic and abiotic stress in *Solanum lycopersicum* (Heinz).** A total of 19 transcriptomic datasets representing various biotic (Bacterial, fungal, viral, and insects) and abiotic stress (Heat, drought, salt, endoplasmic reticulum, cold, and ABA) were analyzed. When log2(FC) values were not available, differential expression analysis was performed using DESeq2. Lectin gene expression was considered significant based on a threshold of log2(FC) > 1 and p < 0.05.

**Table S8: Intra-genomic collinearity analysis of lectin genes in *Solanum* genomes.** MCScanX was performed to identify collinear lectin gene pairs in *S. lycopersicum* (Heinz), *S. lycopersicoides*, *S. pimpinellifolium*, and *S. lycopersicum var. cerasiforme*.

**Table S9: Summary statistics of orthogroup comparisons between *Vitis vinifera* and *Solanum* genomes.** This table presents overall statistics derived from OrthoFinder analysis, including the total number of orthogroups identified per species and the subset corresponding to lectin-specific orthogroups.

**Table S10: Specific orthogroups containing lectin genes in *Vitis vinifera* and *Solanum* genomes.** This table lists orthogroup IDs that uniquely contain lectin genes in each species, along with the corresponding gene identifiers, highlighting species-specific expansions or unique retention of lectin orthologs.

**Table S11: Screening of lectin gene orthogroups for significant expansion and contraction events.** This table presents results from CAFE5 analysis, identifying lectin-containing orthogroups that exhibit significant expansion or contraction (p < 0.05) across *Vitis vinifera* and *Solanum* genomes, based on gene family evolution rates.

**Table S12: Cross-species Ka/Ks analysis of expanded lectin genes in *Solanum lycopersicum* (Heinz).** This table presents non-synonymous (Ka) and synonymous (Ks) substitution rate calculations for lectin genes in significantly expanded orthogroups (p < 0.05), using Ka/Ks Calculator 2.0. Each S. lycopersicum gene is compared to its putative ancestral orthologue in *V. vinifera* (for ancestral expansions) or *S. lycopersicoides* (for lineage-specific expansions) to assess selective evolutionary pressures.

**Table S13: Expression-based functional diversification of paralogues within expanded orthogroups in *S. lycopersicum* (Heinz).** This table presents paralogue pairs identified within each expanded orthogroup, along with their duplication type, Ka/Ks values, expression profiles across various developmental tissues, and co-expression metrics. Based on expression divergence and co-expression patterns, the probable modes of functional evolution for each paralogue pair are inferred.

**Table S14: Validation of expression-based functional diversification of *S. lycopersicum* (Heinz) paralogue pairs in *S. macrocarpon*.** This table presents orthologues of S. lycopersicum paralogue pairs identified in *S. macrocarpon* using reciprocal blast analysis along with expression profiles across various developmental tissues, and co-expression metrics. Based on expression divergence and co-expression patterns, the probable modes of functional evolution for each paralogue pair were validated in *S. macrocarpon*.

**Table S15: Validation of expression-based functional diversification of *S. lycopersicum* (Heinz) paralogue pairs in *S. candidum*.** This table presents orthologues of S. lycopersicum paralogue pairs identified in *S. candidum* using reciprocal blast analysis along with expression profiles across various developmental tissues, and co-expression metrics. Based on expression divergence and co-expression patterns, the probable modes of functional evolution for each paralogue pair were validated in *S. candidum*.

**Table S16: Validation of expression-based functional diversification of *S. lycopersicum* (Heinz) paralogue pairs in *S. quitoense*.** This table presents orthologues of S. lycopersicum paralogue pairs identified in *S. quitoense* using reciprocal blast analysis along with expression profiles across various developmental tissues, and co-expression metrics. Based on expression divergence and co-expression patterns, the probable modes of functional evolution for each paralogue pair were validated in *S. quitoense*.

**Table S17: Validation of expression-based functional diversification of *S. lycopersicum* (Heinz) paralogue pairs in *S. prinophyllum*.** This table presents orthologues of *S. lycopersicum* paralogue pairs identified in *S. prinophyllum* using reciprocal blast analysis along with expression profiles across various developmental tissues, and co-expression metrics. Based on expression divergence and co-expression patterns, the probable modes of functional evolution for each paralogue pair were validated in *S. prinophyllum*.

**Table S18: Primers used in the study.** This table contains forward and reverse primer sequences used to perform RT-qPCR and VIGS analysis.

## Notes

https://github.com/ERR-lab/Lectin-gene-family-in-tomato

### Competing Interest Statement

The authors have declared no competing interest.

## References

An, Z., Yang, Z., Zhou, Y., Huo, S., Zhang, S., Wu, D. et al. (2024). OsJRL negatively regulates rice cold tolerance via interfering phenylalanine metabolism and flavonoid biosynthesis. Plant, Cell & Environment, 47, 4071–4085.

Benoit, M., Jenike, K. M., Satterlee, J. W., Ramakrishnan, S., Gentile, I., Hendelman, A., Passalacqua, M. J., Suresh, H., Shohat, H., Robitaille, G. M., Fitzgerald, B., Alonge, M., Wang, X., Santos, R., He, J., Ou, S., Golan, H., Green, Y., Swartwood, K., … Lippman, Z. B. (2025). Solanum pan-genetics reveals paralogues as contingencies in crop engineering. Nature, 640(8057), 135–145.

Birchler, J. A., & Veitia, R. A. (2014). The Gene Balance Hypothesis: dosage effects in plants. Methods in Molecular Biology (Clifton, N.J.), 1112, 25–32.

Caldwell, D., Kim, B. S., & Iyer-Pascuzzi, A. S. (2017). *Ralstonia solanacearum* differentially colonizes roots of resistant and susceptible tomato plants. Phytopathology, 107(5), 528–536.

Capella-Gutiérrez, S., Silla-Martínez, J. M., & Gabaldón, T. (2009). trimAl: a tool for automated alignment trimming in large-scale phylogenetic analyses. Bioinformatics (Oxford, England), 25(15), 1972–1973.

Catanzariti, A.-M., Lim, G.T.T. and Jones, D.A. (2015), The tomato *I-3* gene: a novel gene for resistance to Fusarium wilt disease. New Phytologist, 207: 106–118.

Castañeda, L., Giménez, E., Pineda, B., García-Sogo, B., Ortiz-Atienza, A., Micol-Ponce, R., Angosto, T., Capel, J., Moreno, V., Yuste-Lisbona, F. J., & Lozano, R. (2022). Tomato CRABS CLAW paralogues interact with chromatin remodelling factors to mediate carpel development and floral determinacy. The New Phytologist, 234(3), 1059–1074.

Chandan, R. K., Kumar, R., Swain, D. M., Ghosh, S., Bhagat, P. K., Patel, S., … & Jha, G. (2023). RAV1 family members function as transcriptional regulators and play a positive role in plant disease resistance. The Plant Journal, 114(1), 39–54.

Chen, N., Shao, Q., Lu, Q., Li, X., & Gao, Y. (2022). Transcriptome analysis reveals differential transcription in tomato (*Solanum lycopersicum*) following inoculation with *Ralstonia solanacearum*. Scientific Reports, 12(1), 22137.

Chen X, Shang J, Chen D, Lei C, Zou Y, Zhai W, Liu G, Xu J, Ling Z, Cao G et al. (2006). A B-lectin receptor kinase gene conferring rice blast resistance. Plant Journal 46: 794–804.

Clark, J. W. (2023). Genome evolution in plants and the origins of innovation. New Phytologist, 240(6), 2204–2209.

Conant, G. C., & Wolfe, K. H. (2008). Turning a hobby into a job: How duplicated genes find new functions. Nature Reviews Genetics, 9(12), 938–950.

Conant, G. C., Birchler, J. A., & Pires, J. C. (2014). Dosage, duplication, and diploidization: clarifying the interplay of multiple models for duplicate gene evolution over time. Current Opinion in Plant Biology, 19, 91–98.

Dang, L., & Van Damme, E. J. M. (2016). Genome-wide identification and domain organization of lectin domains in cucumber. Plant Physiology and Biochemistry, 108, 165–176.

Dang, L., Rougé, P., & Van Damme, E. J. M. (2017). Amaranthin-Like Proteins with Aerolysin Domains in Plants. Frontiers in Plant Science, Volume 8-.

De Bodt, S., Maere, S., & Van de Peer, Y. (2005). Genome duplication and the origin of angiosperms. Trends in Ecology & Evolution, 20(11), 591–597.

De, K., Pal, D., Shanks, C. M., Yates, T. B., Feng, K., Jawdy, S. S., … & Muchero, W. (2023). The Plasminogen-Apple-Nematode (PAN) domain suppresses JA/ET defense pathways in plants. bioRxiv, 2023-06.

De Punder, K., & Pruimboom, L. (2013). The Dietary Intake of Wheat and other Cereal Grains and Their Role in Inflammation. Nutrients, 5(3), 771–787.

Delorme, V., Giranton, J. L., Hatzfeld, Y., Friry, A., Heizmann, P., Ariza, M. J., … & Cock, J. M. (1995). Characterization of the S locus genes, SLG and SRK, of the Brassica S3 haplotype: identification of a membrane-localized protein encoded by the S locus receptor kinase gene. The Plant Journal, 7(3), 429–440.

Domazet-Lošo, M., Široki, T., Šimičević, K. et al. (2024). Macroevolutionary dynamics of gene family gain and loss along multicellular eukaryotic lineages. Nature Communication, 15, 2663.

Eggermont, L., Verstraeten, B., & Van Damme, E. J. M. (2017). Genome-Wide Screening for Lectin Motifs in Arabidopsis thaliana. The Plant Genome, 10(2), plantgenome2017.02.0010.

Emms, D. M., & Kelly, S. (2019). OrthoFinder: phylogenetic orthology inference for comparative genomics. Genome Biology, 20(1), 238.

Esch, L., Kirsch, C., Vogel, L., Kelm, J., Huwa, N., Schmitz, M., … & Schaffrath, U. (2022). Pathogen resistance depending on jacalin-dirigent chimeric proteins is common among Poaceae but absent in the dicot Arabidopsis as evidenced by analysis of homologous single-domain proteins. Plants, 12(1), 67.

Ezoe, A., Shirai, K., & Hanada, K. (2020). Degree of Functional Divergence in Duplicates Is Associated with Distinct Roles in Plant Evolution. Molecular Biology and Evolution, 38(4), 1447–1459.

Ahmed, F., Dola, F. S., Zohra, F. T., Rahman, S. M., Konak, J. N., & Sarkar, M. A. R. (2023). Genome-wide identification, classification, and characterization of lectin gene superfamily in sweet orange (Citrus sinensis L.). PloS One, 18(11), e0294233.

Forslund, K., Henricson, A., Hollich, V., & Sonnhammer, E. L. L. (2008). Domain Tree-Based Analysis of Protein Architecture Evolution. Molecular Biology and Evolution, 25(2), 254–264.

Hamilton, C. D., Zaricor, B., Dye, C. J., Dresserl, E., Michaels, R., & Allen, C. (2024). Ralstonia solanacearum pandemic lineage strain UW551 overcomes inhibitory xylem chemistry to break tomato bacterial wilt resistance. Molecular plant pathology, 25(1), e13395.

He, X., Li, L., Xu, H., Xi, J., Cao, X., Xu, H.,… & Xu, Z. (2017). A rice jacalin-related mannose-binding lectin gene, OsJRL, enhances *Escherichia coli* viability under high salinity stress and improves salinity tolerance of rice. Plant Biology, 19, 257–267.

Hiles, R., Rogers, A., Jaiswal, N., Zhang, W., Butchacas, J., Merfa, M. V., … & Iyer-Pascuzzi, A. S. (2024). A Ralstonia solanacearum type III effector alters the actin and microtubule cytoskeleton to promote bacterial virulence in plants. PLoS Pathogens, 20(12), e1012814.

Hofberger, J. A., Nsibo, D. L., Govers, F., Bouwmeester, K., & Schranz, M. E. (2015). A Complex Interplay of Tandem- and Whole-Genome Duplication Drives Expansion of the L-Type Lectin Receptor Kinase Gene Family in the Brassicaceae. Genome Biology and Evolution, 7(3), 720–734.

Huang, Y., Zhang, L., Zhang, K., Chen, S., Hu, J., & Cheng, F. (2022). The impact of tandem duplication on gene evolution in Solanaceae species. Journal of Integrative Agriculture, 21(4), 1004–1014.

Ingel, B., Caldwell, D., Duong, F., Parkinson, D. Y., McCulloh, K. A., Iyer-Pascuzzi, A. S., … & Lowe-Power, T. M. (2022). Revisiting the source of wilt symptoms: X-ray microcomputed tomography provides direct evidence that Ralstonia biomass clogs xylem vessels. PhytoFrontiers™, 2(1), 41–51.

Jaillon, O., Aury, J.-M., Noel, B., Policriti, A., Clepet, C., Casagrande, A., Choisne, N., Aubourg, S., Vitulo, N., Jubin, C., Vezzi, A., Legeai, F., Hugueney, P., Dasilva, C., Horner, D., Mica, E., Jublot, D., Poulain, J., Bruyère, C., … Characterization, T. F. P. C. for G. G. (2007). The grapevine genome sequence suggests ancestral hexaploidization in major angiosperm phyla. Nature, 449(7161), 463–467.

Katoh, K., Rozewicki, J., & Yamada, K. D. (2019). MAFFT online service: multiple sequence alignment, interactive sequence choice and visualization. Briefings in Bioinformatics, 20(4), 1160–1166.

Kim, N. H., Lee, D. H., Choi, D. S., & Hwang, B. K. (2015). The pepper GNA-related lectin and PAN domain protein gene, CaGLP1, is required for plant cell death and defense signaling during bacterial infection. Plant Science, 241, 307–315.

Kodama, K., Rich, M. K., Yoda, A., Shimazaki, S., Xie, X., Akiyama, K., Mizuno, Y., Komatsu, A., Luo, Y., Suzuki, H., Kameoka, H., Libourel, C., Keller, J., Sakakibara, K., Nishiyama, T., Nakagawa, T., Mashiguchi, K., Uchida, K., Yoneyama, K., … Kyozuka, J. (2022). An ancestral function of strigolactones as symbiotic rhizosphere signals. Nature Communications, 13(1), 3974.

Kumar, R., Barman, A., Phukan, T., Kabyashree, K., Singh, N., Jha, G., … & Ray, S. K. (2017). Ralstonia solanacearum virulence in tomato seedlings inoculated by leaf clipping. Plant Pathology, 66(5), 835–841.

Kwon, C.-T., Tang, L., Wang, X., Gentile, I., Hendelman, A., Robitaille, G., Van Eck, J., Xu, C., & Lippman, Z. B. (2022). Dynamic evolution of small signalling peptide compensation in plant stem cell control. Nature Plants, 8(4), 346–355.

Labbé, J., Muchero, W., Czarnecki, O., Wang, J., Wang, X., Bryan, A. C., … & Tuskan, G. A. (2019). Mediation of plant–mycorrhizal interaction by a lectin receptor-like kinase. Nature Plants, 5(7), 676–680.

Lannoo, N., & Van Damme, E. J. M. (2014). Lectin domains at the frontiers of plant defense. Frontiers in Plant Science, 5.

Li, Z., Defoort, J., Tasdighian, S., Maere, S., Van de Peer, Y., & De Smet, R. (2016). Gene Duplicability of Core Genes Is Highly Consistent across All Angiosperms. The Plant Cell, 28(2), 326–344.

Luo, X., Xu, N., Huang, J., Gao, F., Zou, H., Boudsocq, M., … & Liu, J. (2017). A lectin receptor-like kinase mediates pattern-triggered salicylic acid signalling. Plant Physiology, 174(4), 2501–2514.

Ma, L., Haile, Z. M., Sabbadini, S., Mezzetti, B., Negrini, F., & Baraldi, E. (2022). Functional characterization of MANNOSE-BINDING LECTIN 1, a G-type lectin gene family member, in response to fungal pathogens of strawberry. Journal of Experimental Botany, 74(1), 149–161.

Maere, S., De Bodt, S., Raes, J., Casneuf, T., Van Montagu, M., Kuiper, M., & Van de Peer, Y. (2005). Modeling gene and genome duplications in eukaryotes. Proceedings of the National Academy of Sciences of the United States of America, 102(15), 5454–5459.

Maimbo, M., Ohnishi, K., Hikichi, Y., Yoshioka, H., & Kiba, A. (2010). S-glycoprotein-like protein regulates defense responses in Nicotiana plants against Ralstonia solanacearum. Plant Physiology, 152(4), 2023–2035.

McLeay, R. C., & Bailey, T. L. (2010). Motif Enrichment Analysis: a unified framework and an evaluation on ChIP data. BMC Bioinformatics, 11(1), 165.

Mendes, F. K., Vanderpool, D., Fulton, B., & Hahn, M. W. (2021). CAFE 5 models variation in evolutionary rates among gene families. Bioinformatics (Oxford, England), 36(22–23), 5516– 5518.

Micol-Ponce, R., García-Alcázar, M., Lebrón, R., Capel, C., Pineda, B., García-Sogo, B., Alché, J. de D., Ortiz-Atienza, A., Bretones, S., Yuste-Lisbona, F. J., Moreno, V., Capel, J., & Lozano, R. (2023). Tomato POLLEN DEFICIENT 2 encodes a G-type lectin receptor kinase required for viable pollen grain formation. Journal of Experimental Botany, 74(1), 178–193.

Minh, B. Q., Schmidt, H. A., Chernomor, O., Schrempf, D., Woodhams, M. D., von Haeseler, A., & Lanfear, R. (2020). IQ-TREE 2: New Models and Efficient Methods for Phylogenetic Inference in the Genomic Era. Molecular Biology and Evolution, 37(5), 1530–1534.

Mishra, A., Behura, A., Mawatwal, S., Kumar, A., Naik, L., Mohanty, S. S., & others. (2019). Structure-function and application of plant lectins in disease biology and immunity. Food and Chemical Toxicology, 134, 110827.

Naithani, S., Komath, S. S., Nonomura, A., & Govindjee, G. (2021). Plant lectins and their many roles: Carbohydrate-binding and beyond. Journal of Plant Physiology, 266, 153531.

Nonomura, A. M., & Benson, A. A. (2014). The path of carbon in photosynthensis. XXXI. The role of lectins. Journal of Plant Nutrition, 37(6), 785–794.

Ohno S. Evolution by gene duplication. Berlin (Germany): Springer-Verlag; 1970.

Osman, M. E. M., Osman, R. S. H., Elmubarak, S. A. A., Ibrahim, M. A., Abakar, H. B. M., Dirar, A. I., & Konozy, E. H. E. (2024). In silico analysis of L- and G-type lectin receptor kinases in tomato: evolution, diversity, and abiotic responses. BMC Genomics, 25(1), 1143.

Pan, J., Li, Z., Wang, Q., Yang, L., Yao, F., & Liu, W. (2020). An S-domain receptor-like kinase, OsESG1, regulates early crown root development and drought resistance in rice. Plant Science, 290, 110318.

Panchy, N., Lehti-Shiu, M., & Shiu, S.-H. (2016). Evolution of Gene Duplication in Plants. Plant Physiology, 171(4), 2294–2316.

Peng, X., Wang, M., Li, Y., Yan, W., Chang, Z., Chen, Z., Xu, C., Yang, C., Deng, X. W., Wu, J., & Tang, X. (2020). Lectin receptor kinase OsLecRK-S.7 is required for pollen development and male fertility. Journal of Integrative Plant Biology, 62(8), 1227–1245.

Petrova, N., Nazipova, A., Gorshkov, O., Mokshina, N., Patova, O., & Gorshkova, T. (2021). Gene Expression Patterns for Proteins With Lectin Domains in Flax Stem Tissues Are Related to Deposition of Distinct Cell Wall Types. Frontiers in Plant Science, Volume 12.

Qiao, Z., Yates, T. B., Shrestha, H. K., Engle, N. L., Flanagan, A., Morrell-Falvey, J. L., … & Chen, J. G. (2021). Towards engineering ectomycorrhization into switchgrass bioenergy crops via a lectin receptor-like kinase. Plant biotechnology journal, 19(12), 2454–2468.

Rairdan, G. J., Collier, S. M., Sacco, M. A., Baldwin, T. T., Boettrich, T., & Moffett, P. (2008). The coiled-coil and nucleotide binding domains of the Potato Rx disease resistance protein function in pathogen recognition and signaling. The Plant Cell, 20(3), 739–751.

Ramireddy, E., Hosseini, S.A., Eggert, K., Gillandt, S., Gnad, H., von Wirén, N., and Schm€ ulling, T. (2018). Root Engineering in Barley: Increasing Cytokinin Degradation Produces a Larger Root System, Mineral Enrichment in the Shoot and Improved Drought Tolerance. Plant Physiol. 177:1078–1095.

Saeed, B., Baranwal, V. K., & Khurana, P. (2016). Identification and Expression Profiling of the Lectin Gene Superfamily in Mulberry. The Plant Genome, 9(2).

Santamaría, M. E., Martínez, M., Arnaiz, A., Rioja, C., Burow, M., Grbic, V., & Díaz, I. (2019). An Arabidopsis TIR-Lectin Two-Domain Protein Confers Defense Properties against Tetranychus urticae. Plant Physiology, 179(4), 1298–1314.

Sato, S., Tabata, S., Hirakawa, H., Asamizu, E., Shirasawa, K., Isobe, S., Kaneko, T., Nakamura, Y., Shibata, D., Aoki, K., Egholm, M., Knight, J., Bogden, R., Li, C., Shuang, Y., Xu, X., Pan, S., Cheng, S., Liu, X., … Fabra, U. P. (2012). The tomato genome sequence provides insights into fleshy fruit evolution. Nature, 485(7400), 635–641.

Shinjo, A., Araki, Y., Hirano, K., Arie, T., Ugaki, M., & Teraoka, T. (2011). Transgenic rice plants that over-express the mannose-binding rice lectin have enhanced resistance to rice blast. Journal of General Plant Pathology, 77, 85–92.

Soltis, P. S., & Soltis, D. E. (2021). Plant genomes: Markers of evolutionary history and drivers of evolutionary change. PLANTS, PEOPLE, PLANET, 3(1), 74–82.

Stefanowicz, K., Lannoo, N., Zhao, Y., Eggermont, L., Van Hove, J., Al Atalah, B., & Van Damme, E. J. M. (2016). Glycan-binding F-box protein from Arabidopsis thaliana protects plants from Pseudomonas syringae infection. BMC Plant Biology, 16(1), 213.

Streubel, S., Deiber, S., Rötzer, J., Mosiolek, M., Jandrasits, K., & Dolan, L. (2023). Meristem dormancy in Marchantia polymorpha is regulated by a liverwort-specific miRNA and a clade III SPL gene. Current Biology : CB, 33(4), 660–674.e4.

The Amborella genome and the evolution of flowering plants. (2013). Science (New York, N.Y.), 342(6165), 1241089.

Trontin, C., Kiani, S., Corwin, J. A., Hématy, K., Yansouni, J., Kliebenstein, D. J., & Loudet, O. (2014). A pair of receptor-like kinases is responsible for natural variation in shoot growth response to mannitol treatment in rabidopsis thaliana. The Plant Journal, 78(1), 121–133.

Tsaneva, M., & Van Damme, E. J. M. (2020). 130 years of Plant Lectin Research. Glycoconjugate Journal, 37(5), 533–551.

Vaattovaara, A., Brandt, B., Rajaraman, S., Safronov, O., Veidenberg, A., Luklová, M., Kangasjärvi, J., Löytynoja, A., Hothorn, M., Salojärvi, J., & Wrzaczek, M. (2019). Mechanistic insights into the evolution of DUF26-containing proteins in land plants. Communications Biology, 2(1), 56.

Van Damme, E. J. (2022). 35 years in plant lectin research: a journey from basic science to applications in agriculture and medicine. Glycoconjugate journal, 39(1), 83–97.

Van Damme, E. J. M., Lannoo, N., & Peumans, W. J. (2008). Plant Lectins (J.-C. Kader & M. B. T.-A. in B. R. Delseny (Eds.); Vol. 48, pp. 107–209). Academic Press.

Van Damme, E. J., Nakamura-Tsuruta, S., Smith, D. F., Ongenaert, M., Winter, H. C., Rougé, P., … & Peumans, W. J. (2007). Phylogenetic and specificity studies of two-domain GNA-related lectins: generation of multispecificity through domain duplication and divergent evolution. Biochemical Journal, 404(1), 51–61.

Van de Peer, Y., Mizrachi, E., & Marchal, K. (2017). The evolutionary significance of polyploidy. Nature Reviews. Genetics, 18(7), 411–424.

Van Holle, S., & Van Damme, E. J. M. (2015). Distribution and Evolution of the Lectin Family in Soybean (Glycine max). Molecules, 20(2), 2868–2891.

Van Holle, S., De Schutter, K., Eggermont, L., Tsaneva, M., Dang, L., & Van Damme, E. J. M. (2017). Comparative Study of Lectin Domains in Model Species: New Insights into Evolutionary Dynamics. International Journal of Molecular Sciences, 18(6), 1136.

Van Holle, S., Rougé, P., & Van Damme, E. J. M. (2017). Evolution and structural diversification of *Nictaba*-like lectin genes in food crops with a focus on soybean (*Glycine max*), Annals of Botany, 119(5), 901–914.

Van Holle, S., & Van Damme, E. J. M. (2019). Messages From the Past: New Insights in Plant Lectin Evolution. Frontiers in Plant Science, 10.

Van Hove, J., De Jaeger, G., De Winne, N., Guisez, Y., & Van Damme, E. J. M. (2015). The Arabidopsis lectin EULS3 is involved in stomatal closure. Plant Science, 238, 312–322.

Wang, D., Zhang, Y., Zhang, Z., Zhu, J., & Yu, J. (2010). KaKs_Calculator 2.0: A Toolkit Incorporating Gamma-Series Methods and Sliding Window Strategies. Genomics, Proteomics & Bioinformatics, 8(1), 77–80.

Wang, J., Chitsaz, F., Derbyshire, M. K., Gonzales, N. R., Gwadz, M., Lu, S., Marchler, G. H., Song, J. S., Thanki, N., Yamashita, R. A., Yang, M., Zhang, D., Zheng, C., Lanczycki, C. J., & Marchler-Bauer, A. (2023). The conserved domain database in 2023. Nucleic Acids Research, 51(D1), D384–D388.

Wang, Y., Bouwmeester, K., Beseh, P., Shan, W., & Govers, F. (2014). Phenotypic Analyses of Arabidopsis T-DNA Insertion Lines and Expression Profiling Reveal That Multiple L-Type Lectin Receptor Kinases Are Involved in Plant Immunity. Molecular Plant-Microbe Interactions®, 27(12), 1390–1402.

Wang, Y., Tang, H., Wang, X., Sun, Y., Joseph, P. V, & Paterson, A. H. (2024). Detection of colinear blocks and synteny and evolutionary analyses based on utilization of MCScanX. Nature Protocols, 19(7), 2206–2229.

Weidenbach, D., Esch, L., Möller, C., Hensel, G., Kumlehn, J., Höfle, C., … & Schaffrath, U. (2016). Polarised defence against fungal pathogens is mediated by the jacalin-related lectin domain of modular Poaceae-specific proteins. Molecular Plant, 9(4), 514–527.

Willmann, R., Lajunen, H. M., Erbs, G., Newman, M.-A., Kolb, D., Tsuda, K., Katagiri, F., Fliegmann, J., Bono, J.-J., Cullimore, J. V, Jehle, A. K., Götz, F., Kulik, A., Molinaro, A., Lipka, V., Gust, A. A., & Nürnberger, T. (2011). Arabidopsis lysin-motif proteins LYM1 LYM3 CERK1 mediate bacterial peptidoglycan sensing and immunity to bacterial infection. Proceedings of the National Academy of Sciences, 108(49), 19824–19829.

Xiao, J., Li, C., Xu, S., Xing, L., Xu, Y., & Chong, K. (2015). JACALIN-LECTIN LIKE1 regulates the nuclear accumulation of GLYCINE-RICH RNA-BINDING PROTEIN7, influencing the RNA processing of FLOWERING LOCUS C antisense transcripts and flowering time in Arabidopsis. Plant physiology, 169(3), 2102–2117.

Xin, Z., Wang, A., Yang, G., Gao, P., & Zheng, Z.-L. (2009). The Arabidopsis A4 Subfamily of Lectin Receptor Kinases Negatively Regulates Abscisic Acid Response in Seed Germination . Plant Physiology, 149(1), 434–444.

Xing, S., Li, M., & Liu, P. (2013). Evolution of S-domain receptor-like kinases in land plants and origination of S-locus receptor kinases in Brassicaceae. BMC Evolutionary Biology, 13(1), 69.

Yamaji, Y., Maejima, K., Komatsu, K., Shiraishi, T., Okano, Y., Himeno, M., … & Namba, S. (2012). Lectin-mediated resistance impairs plant virus infection at the cellular level. The Plant Cell, 24(2), 778–793.

Yanai, I., Benjamin, H., Shmoish, M., Chalifa-Caspi, V., Shklar, M., Ophir, R., Bar-Even, A., Horn-Saban, S., Safran, M., Domany, E., Lancet, D., & Shmueli, O. (2005). Genome-wide midrange transcription profiles reveal expression level relationships in human tissue specification. Bioinformatics, 21(5), 650–659.

Yang, M., Wang, Y., Chen, C., Xin, X., Dai, S., Meng, C., & Ma, N. (2024). Transcription factor WRKY75 maintains auxin homeostasis to promote tomato defense against Pseudomonas syringae. Plant physiology, 195(2), 1053–1068.

Yu, G., Zhang, L., Wang, K., & Macho, A. P. (2023). Inoculation of Arabidopsis seedlings with Ralstonia solanacearum in sterile agar plates. STAR protocols, 4(3), 102474.

Yuan, L., Wu, M., Tan, D., Zhang, S., Zhang, H., Li, J., Xia, G., & Wang, F. (2025). Mannose-binding lectin 1.1A interacts with hypersensitive-induced response 4 to promote hypersensitive cell death and defense responses in cotton upon Verticillium dahliae infection. The Plant Journal, 121(4), e70018.

Zhou, D., Godinez-Vidal, D., He, J., Teixeira, M., Guo, J., Wei, L., Van Norman, J. M., & Kaloshian, I. (2023). A G-type lectin receptor kinase negatively regulates Arabidopsis immunity against root-knot nematodes. Plant Physiology, 193(1), 721–735.

Zipfel, C., & Oldroyd, G. E. D. (2017). Plant signalling in symbiosis and immunity. Nature, 543(7645), 328–336.

