## Supplementary figures and images for "Systematic analysis of lectin gene family reveals dynamic modes of paralogue evolution and immune regulatory functions in tomato"

### Figure_S1

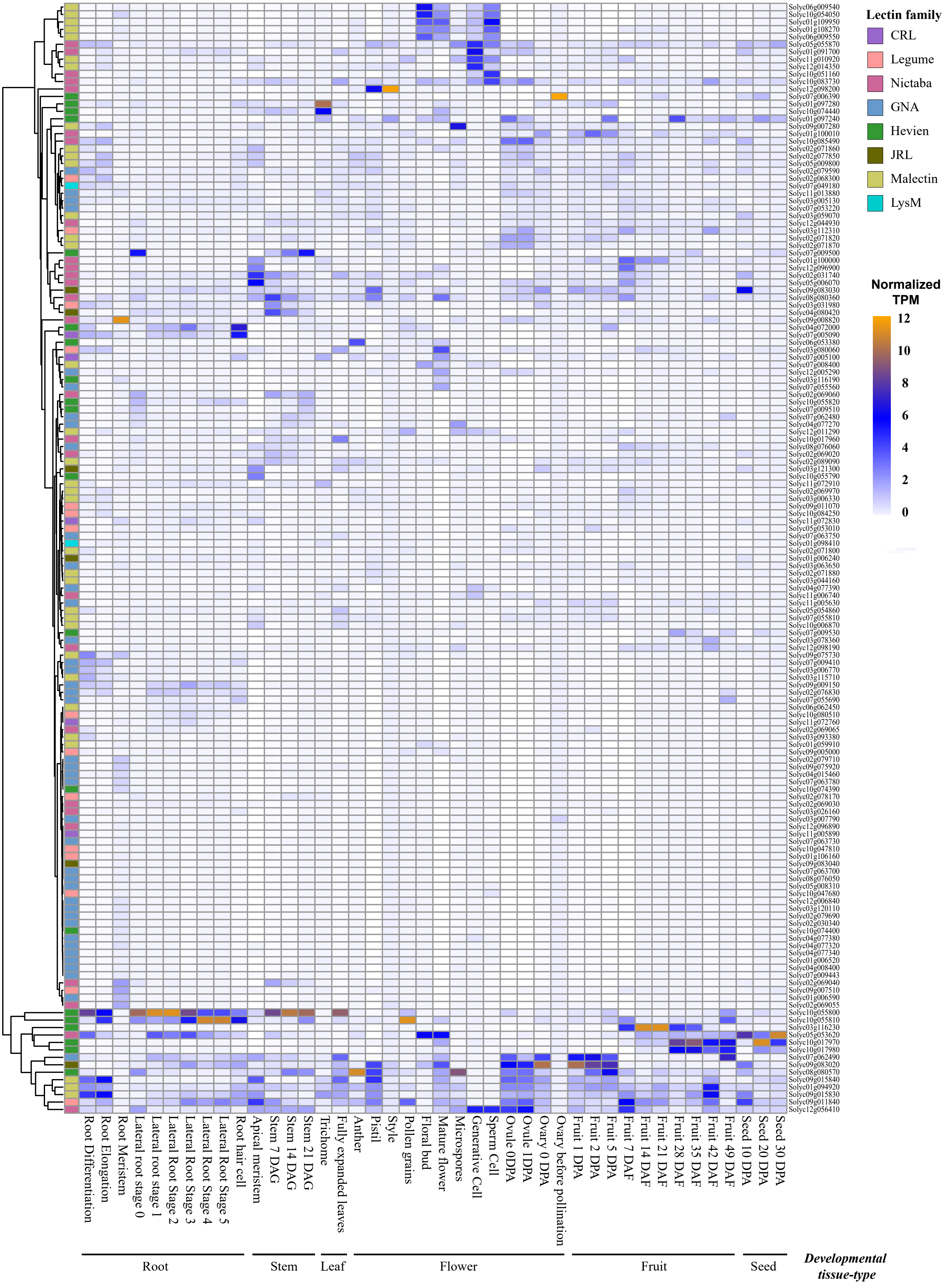


**Figure S1: Transcript abundance of lectin genes in *S. lycopersicum* (Heinz) during development.**

### Figure_S2

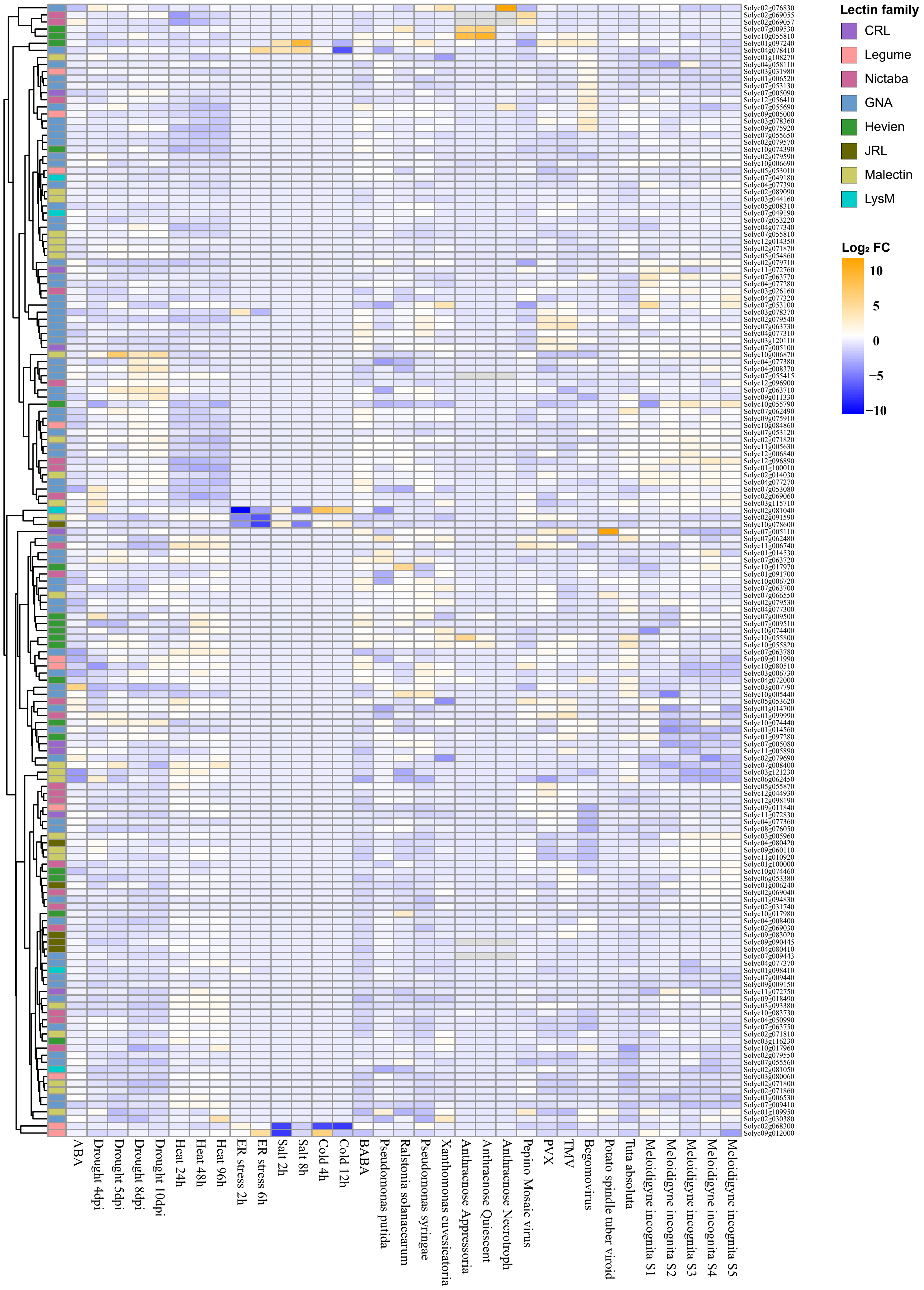


**Figure S2: Transcript abundance of lectin genes in *S. lycopersicum* (Heinz) during stress.**
