## Supplementary material for "Systematic analysis of lectin gene family reveals dynamic modes of paralogue evolution and immune regulatory functions in tomato": Figure_S3

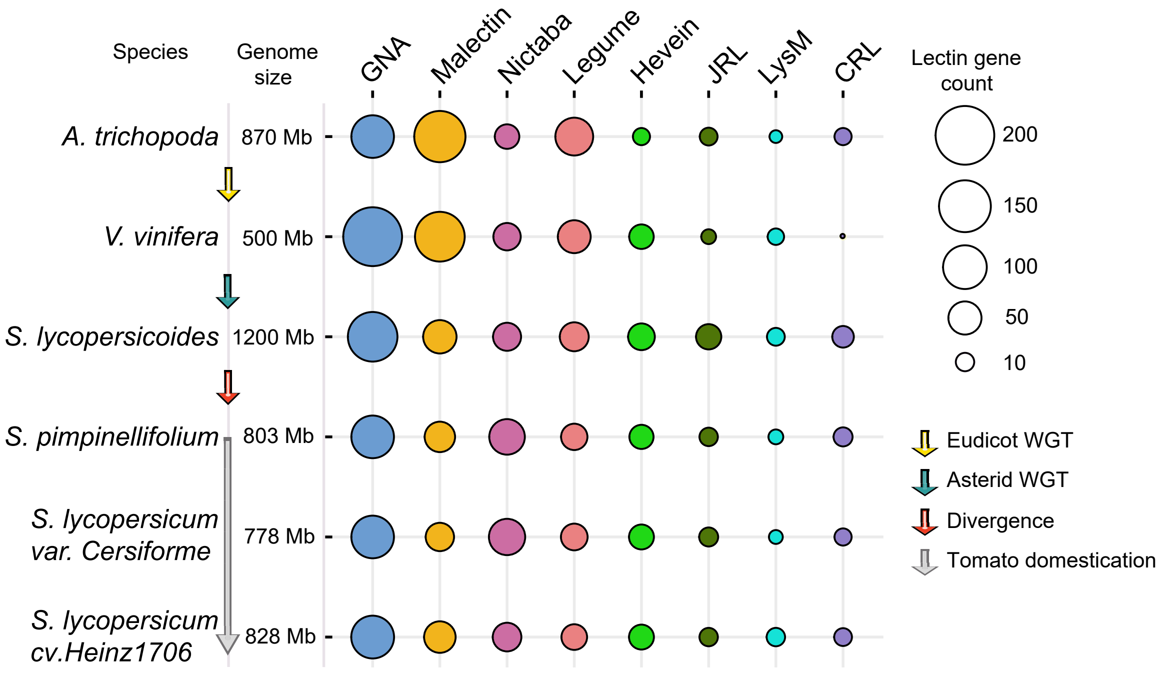


**Figure S3. Comparative distribution of lectin gene counts in Solanum species and their ancestral genomes.**
