## Supplementary material for "Systematic analysis of lectin gene family reveals dynamic modes of paralogue evolution and immune regulatory functions in tomato": Figure_S4

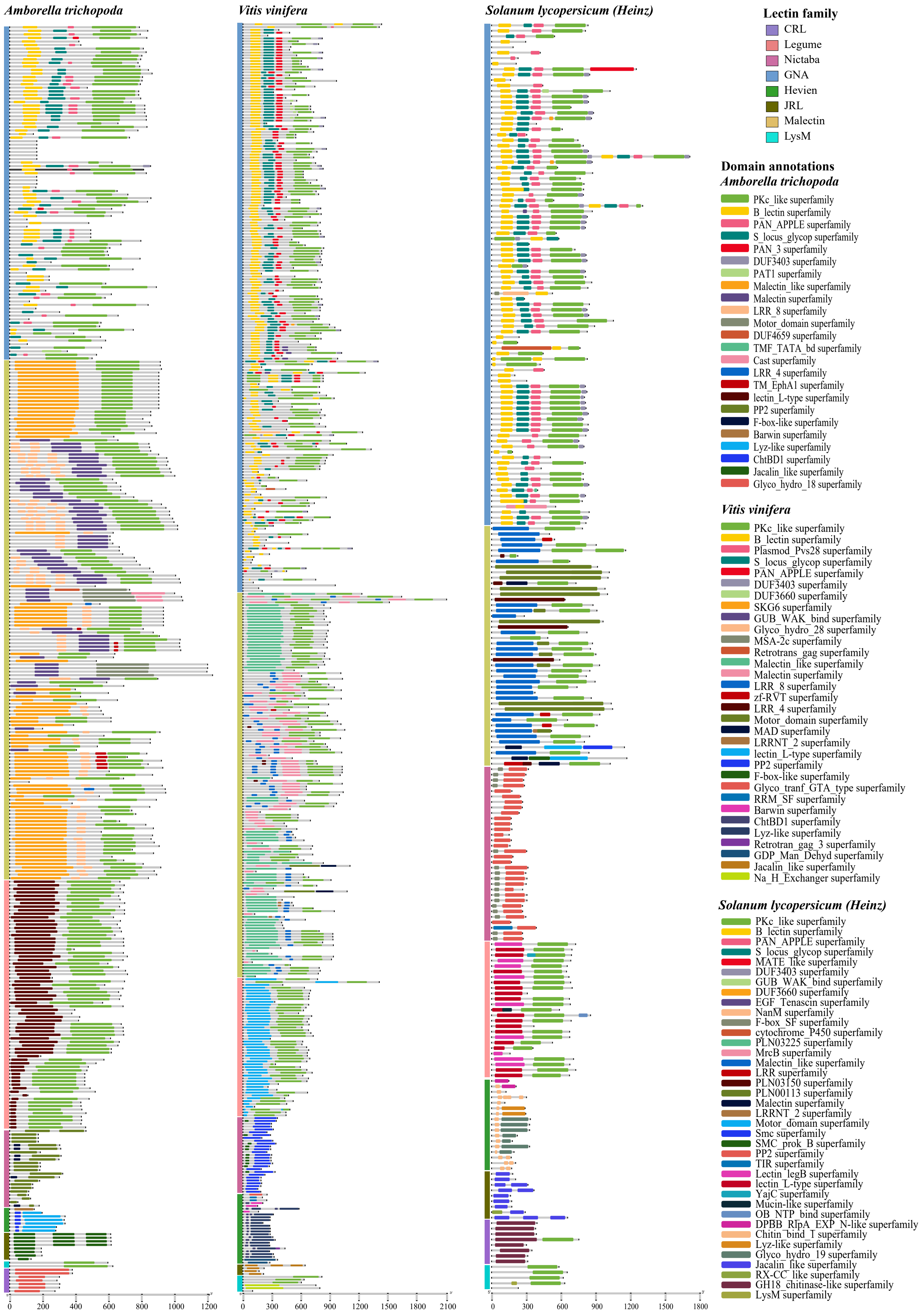


**Figure S4: Protein domain architecture of lectin genes in *S. lycopersicum*, *V. vinifera* and *A. trichopoda*.**
