## Supplementary material for "Systematic analysis of lectin gene family reveals dynamic modes of paralogue evolution and immune regulatory functions in tomato": Figure_S5

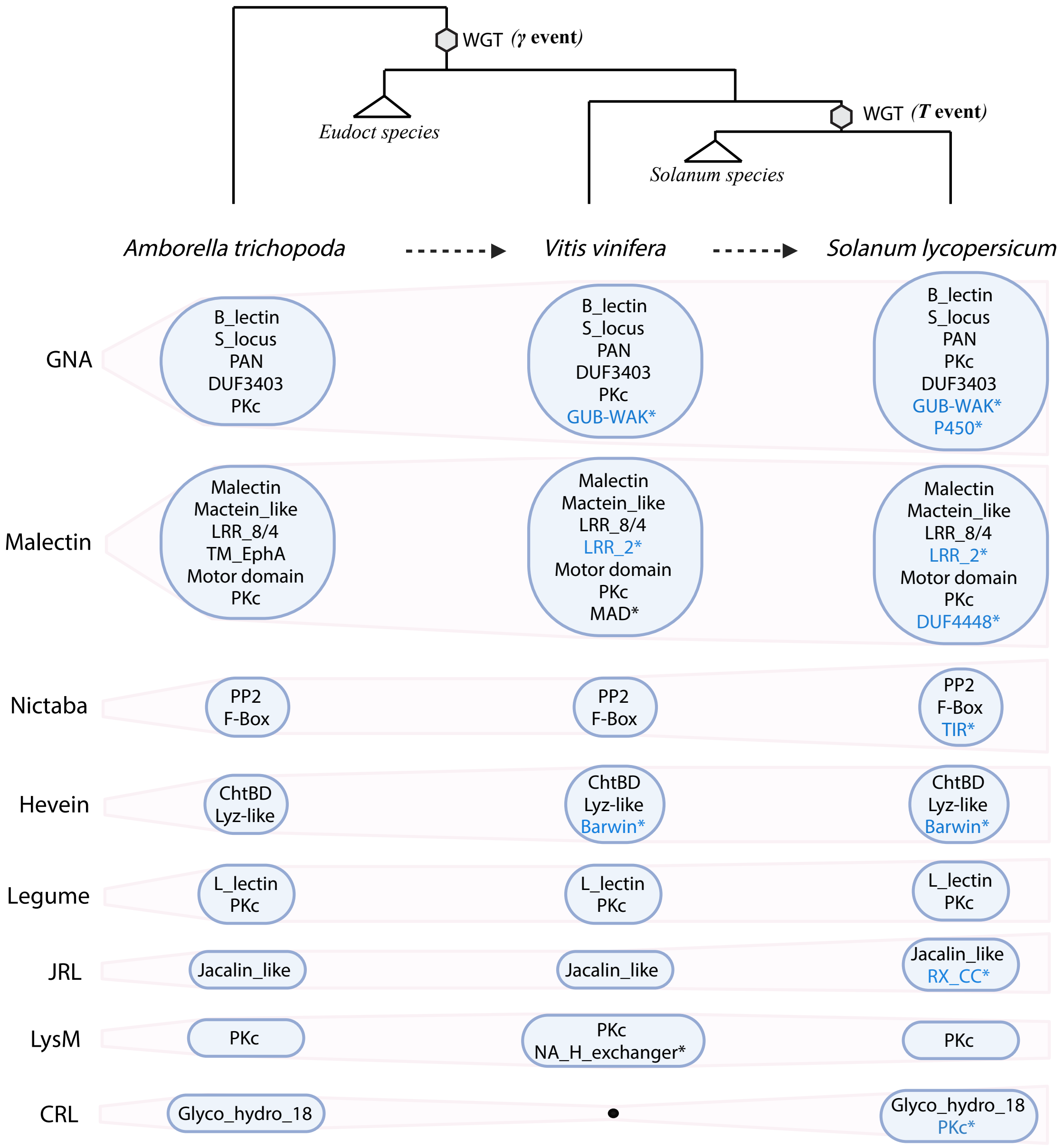


**Figure S5. Impact of whole-genome triplication and duplication events on protein domain expansions in lectin gene families.**
