## Supplementary material for "Systematic analysis of lectin gene family reveals dynamic modes of paralogue evolution and immune regulatory functions in tomato": Figure_S6

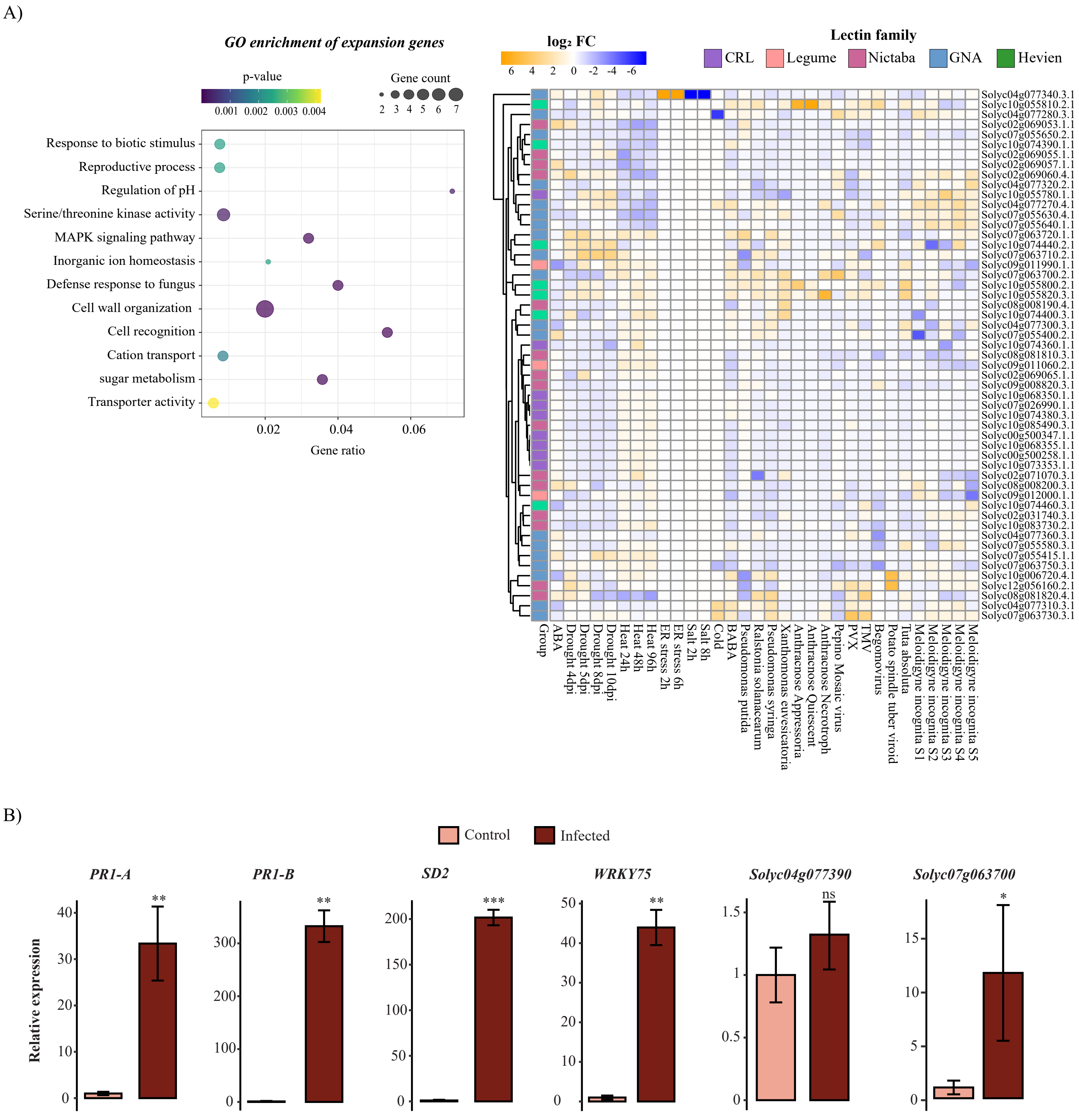


**Figure S6. Functional roles of expanded lectin genes in *S. lycopersicum* (Heinz) under stress conditions.**
