## Supplementary material for "Systematic analysis of lectin gene family reveals dynamic modes of paralogue evolution and immune regulatory functions in tomato": Figure_S7

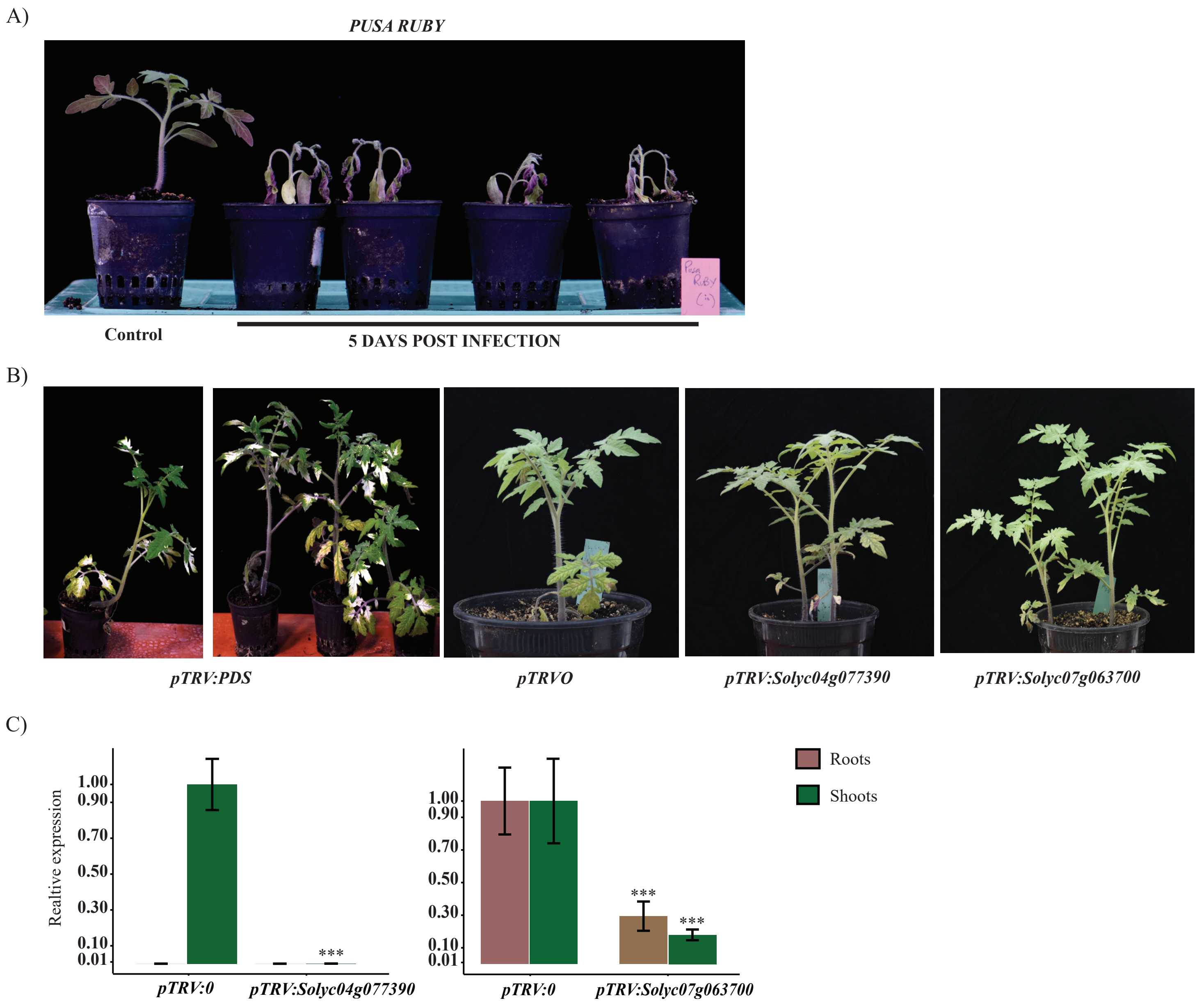


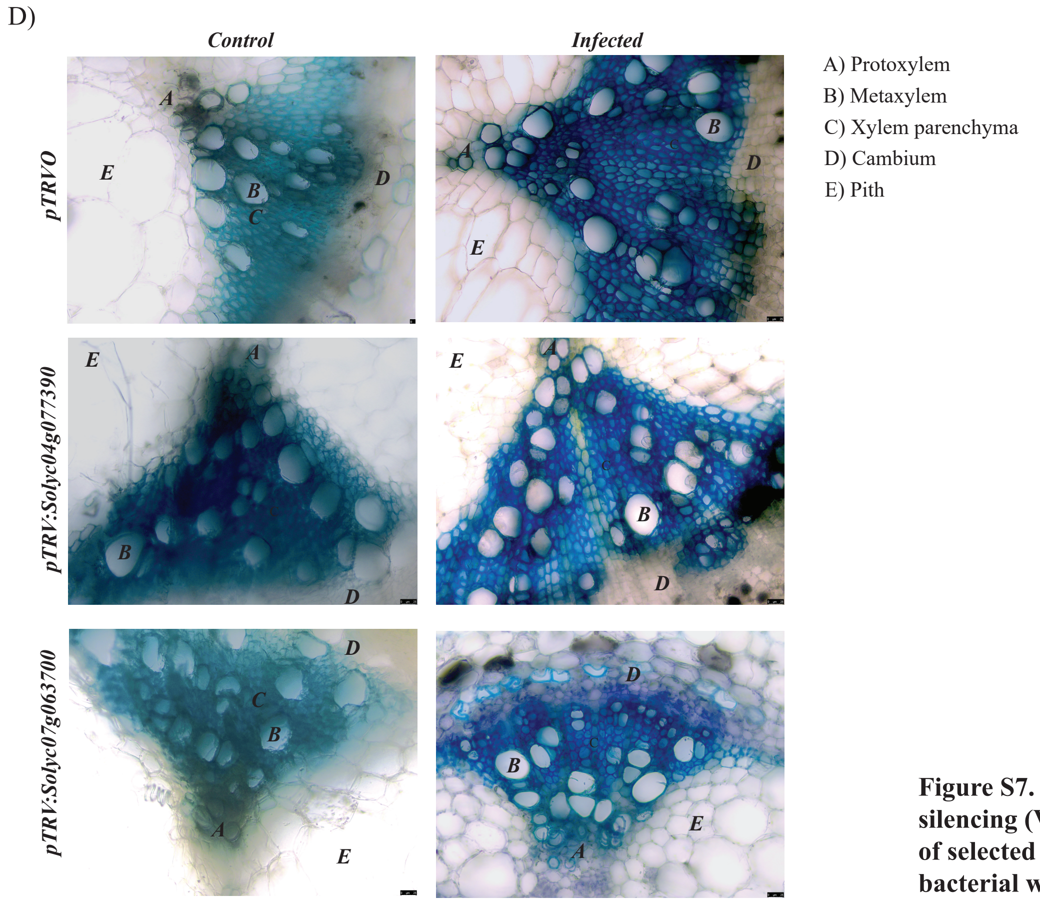


**Figure S7. Assessment of virus-induced gene silencing (VIGS) controls for functional validation of selected lectin genes in *S. lycopersicum* during bacterial wilt infection.**
